# Virus glycoprotein nanodisc platform for vaccine design

**DOI:** 10.1101/2025.05.02.651272

**Authors:** Kimmo Rantalainen, Alessia Liguori, Gabriel Ozorowski, Claudia Flynn, Jon M. Steichen, Olivia Swanson, Patrick J. Madden, Sabyasachi Baboo, Swastik Phulera, Anant Gharpure, Danny Lu, Oleksandr Kalyuzhniy, Patrick Skog, Sierra Terada, Monolina Shil, Jolene K. Diedrich, Erik Georgeson, Ryan Tingle, Saman Eskandarzadeh, Wen-Hsin Lee, Nushin Alavi, Diana Goodwin, Michael Kubitz, Sonya Amirzehni, Sunny Himansu, Devin Sok, Jeong Hyun Lee, John R. Yates, James C. Paulson, Shane Crotty, Torben Schiffner, Andrew B. Ward, William R. Schief

**Affiliations:** Department of Immunology and Microbiology, The Scripps Research Institute, La Jolla, CA, USA; IAVI Neutralizing Antibody Center, The Scripps Research Institute, La Jolla, CA, USA; Consortium for HIV/AIDS Vaccine Development (CHAVD), The Scripps Research Institute, La Jolla, CA, USA; Department of Integrative Structural and Computational Biology, The Scripps Research Institute, La Jolla, CA 92037, USA; Center for Vaccine Innovation, La Jolla Institute for Immunology, La Jolla, CA 92037, USA; Moderna Inc., Cambridge, MA, 02139, USA; Division of Infectious Diseases, Department of Medicine, University of California San Diego, La Jolla, CA 92037, USA; Institute for Drug Discovery, Leipzig University Medical Faculty, Leipzig, 04103, Germany

## Abstract

Transmembrane glycoproteins of enveloped viruses are the targets of neutralizing antibodies and essential vaccine antigens. mRNA-LNP technology allows in situ production of transmembrane glycoproteins upon immunization, but biophysical characterization of transmembrane antigens and in vitro analysis of post-immunization antibody responses typically rely on soluble proteins. Here, we present a methodological platform for assembling transmembrane glycoprotein vaccine candidates into lipid nanodiscs. We demonstrate the utility of the nanodiscs in HIV membrane proximal external region (MPER)-targeting vaccine development by binding assays using surface plasmon resonance (SPR), ex vivo B cell sorting with fluorescence-activated cell sorting (FACS), and by determining the structure of a prototypical HIV MPER-targeting immunogen nanodisc in complex with three broadly neutralizing antibodies (bnAbs), including the MPER bnAb 10E8, to 3.5 Å by cryogenic electron microscopy (cryo-EM), providing a template for structure-based immunogen design for MPER. Overall, the platform offers a tool for accelerating the development of next-generation viral vaccines.

## Summary

In the past two decades, nanodiscs have emerged as means to encapsulate transmembrane proteins into a stable and more native-like environment. In most cases, proteins are extracted from the membrane by detergent solubilization and then assembled into a nanodisc by removal of the detergent in presence of lipid molecules and a scaffold protein. Since the first apolipoprotein-derived membrane scaffold proteins (MSPs) were introduced, several variations and alternative scaffolds have been described, allowing for a variety of disc sizes and versatile experimental approaches^1–5^. Structural studies of membrane proteins have perhaps benefitted the most from nanodiscs, resulting in important advances in understanding of membrane protein structure-function relationship^6, 7^. They have also been used successfully in a variety of functional and biophysical approaches such as surface plasmon resonance (SPR)^8–10^.

Viral glycoproteins of enveloped viruses are membrane proteins and essential targets for vaccine development. Structure-based vaccine design was an integral part of the successful COVID-19 vaccine development^11–13^, and a significant portion of the groundwork stemmed from decades of HIV vaccine development^14^. Structure-based methods, combined with rapidly advancing computational protein design approaches and mRNA lipid nanoparticle (mRNA-LNP) technology, are now revolutionizing iterative vaccine development^15, 16^. Immunogen delivery by mRNA-LNP expands the available immunogens to more native-like constructs that may contain the membrane-proximal epitopes such as the recently developed cleavage independent NFL trimers expressing HIV Env^17^ and the MD39.3 gp151 Env trimer used in the HVTN302 clinical trial (NCT05217641), but recombinant transmembrane glycoprotein production for biophysical characterization of the immunogens remains challenging. In most studies, membrane proteins are truncated before the transmembrane domain to obtain soluble ectodomains with higher expression levels and easier handling and applicability in downstream analytical methods. The regions excluded typically include the membrane-proximal external region (MPER), transmembrane domain (TM), and intracellular C-terminal domain (CT). In influenza HA, the anchor region analogous to HIV MPER joins the ectodomain as flexible linkers to three transmembrane helices and allows the trimer to tilt in relation to bilayer surface^18^. Both Env MPER and HA anchor have highly conserved epitopes for neutralizing antibodies (nAbs) making them attractive targets for vaccine development^19–23^. Significant progress in MPER targeted HIV vaccine development has been made recently with peptide-liposome formulations showing induction of heterologous neutralizing antibody B cell lineages in humans, and with mRNA delivered germline-targeting (GT) epitope scaffold efficiently inducing antibody precursors in non-human primates (NHPs) and mice^24, 25^. Analysis of antibodies elicited by these vaccines as well as further immunogen development would benefit from inclusion of the full epitope in recombinantly expressed proteins. For example, recombinant proteins matching the transmembrane immunogens delivered as mRNA are required for accurate immunogen assessment in iterative vaccine design methods, such as SPR and structural studies. Importantly, transmembrane versions of Env may also be required for more accurate representations of the glycan shield, as many neutralizing antibodies bind specifically to glycans, and glycoproteins expressed as transmembrane proteins have been shown to have glycan shield compositions closer to the native glycan shield^26–28^.

Some uses of nanodiscs to support vaccine development have been demonstrated, particularly in efforts to characterize the structures of HIV MPER-targeting HIV antibody epitopes and transmembrane domains of virus glycoproteins in their native membrane bilayer environment^29–33^. The nanodisc assembly platform presented here provides the reproducibility, scalability, and accurate replication of the vaccine candidate properties that enable routine use of transmembrane viral glycoproteins in key methods employed in iterative, rational vaccine design. We demonstrate use of nanodiscs to measure antibody affinities by SPR, as FACS probes to sort antigen-specific B cells from mouse and non-human primate (NHP) immunization experiments, and for solving the cryo-EM structure of a prototype HIV boost vaccine candidate in complex with three broadly neutralizing antibodies (bnAbs) that target distinct sites of vulnerability on the HIV Env surface. We highlight the benefits of the approach by reporting the structure of the entire protein epitope of an HIV MPER-targeting bnAb 10E8. Finally, we show the adaptability of the platform to other glycoproteins by assembling engineered Ebola virus glycoprotein in nanodiscs using identical conditions to HIV Env.

## Results

### Glycoprotein Nanodisc Assembly and Analysis Workflow

We set to establish a workflow that provides material suitable for methods commonly used in rational, iterative vaccine design (Fig. 1). The workflow was set up using the engineered and highly expressing HIV Env clone BG505 MD39.3 gp151^34,35^ that is currently being tested in a phase 1 clinical trial HVTN 302 (<u>NCT05217641</u>). A linker, followed by an HRV3C protease cleavage site and a strep tag was added to the intracellular C-terminus, leading to a nanodisc assembly construct base that will hereon be referred to as Env gp151 ND (Fig. 2a-b). This construct was then used to optimize the throughput and reproducibility of nanodisc production, leading to a standardized workflow (Fig. 2c). In short, transmembrane glycoproteins (GPs) were expressed on the surface of human FreeStyle 293-F cells, followed by detergent solubilization with TX-100-containing buffer. As the transmembrane domain of viral glycoproteins is composed of only three helices – one per protomer – detergent solubilization by TX-100 was not expected to disturb the quaternary structure of the engineered GP trimer as indicated by the binding of structure specific antibodies to extracted GPs^27, 28^. The detergent-solubilized glycoprotein was then assembled into nanodiscs using the MSP1D1 scaffold with or without a biotin tag. While a variety of scaffold proteins are available, we opted to reduce variables and focused on using the standard MSP1D1 scaffold for all purposes. The developed “batch assembly” approach, resembling previously described “on-column”^36^ and “on-bead”^37^ methods, facilitated reproducibility and efficient assembly while GPs remained bound to the matrix. Empty nanodiscs for control experiments were produced by omitting GP in the assembly step and using His-tag of the scaffold protein with NiNTA affinity purification prior to SEC. Utilizing this workflow, up to 12 samples were routinely processed simultaneously in five days from transfection to purified nanodisc. One liter of cells yielded approximately 100 to 700 µg of pure GP nanodiscs, which was sufficient for one to three endpoint assays. For example, a single production batch produced an EBOV GP nanodiscs, an HIV Env nanodisc-Fab complex for cryo-EM structural studies, and eight FACS probes for B cell sorting. Final GP nanodisc preparations were stable for at least three months at +4°C based on absorbance at 280 nm, presence of trimeric GP in nanodisc in negative stain (ns) EM 2D class averages, and activity in SPR analysis (Fig. 3 b).

**Figure 1.**
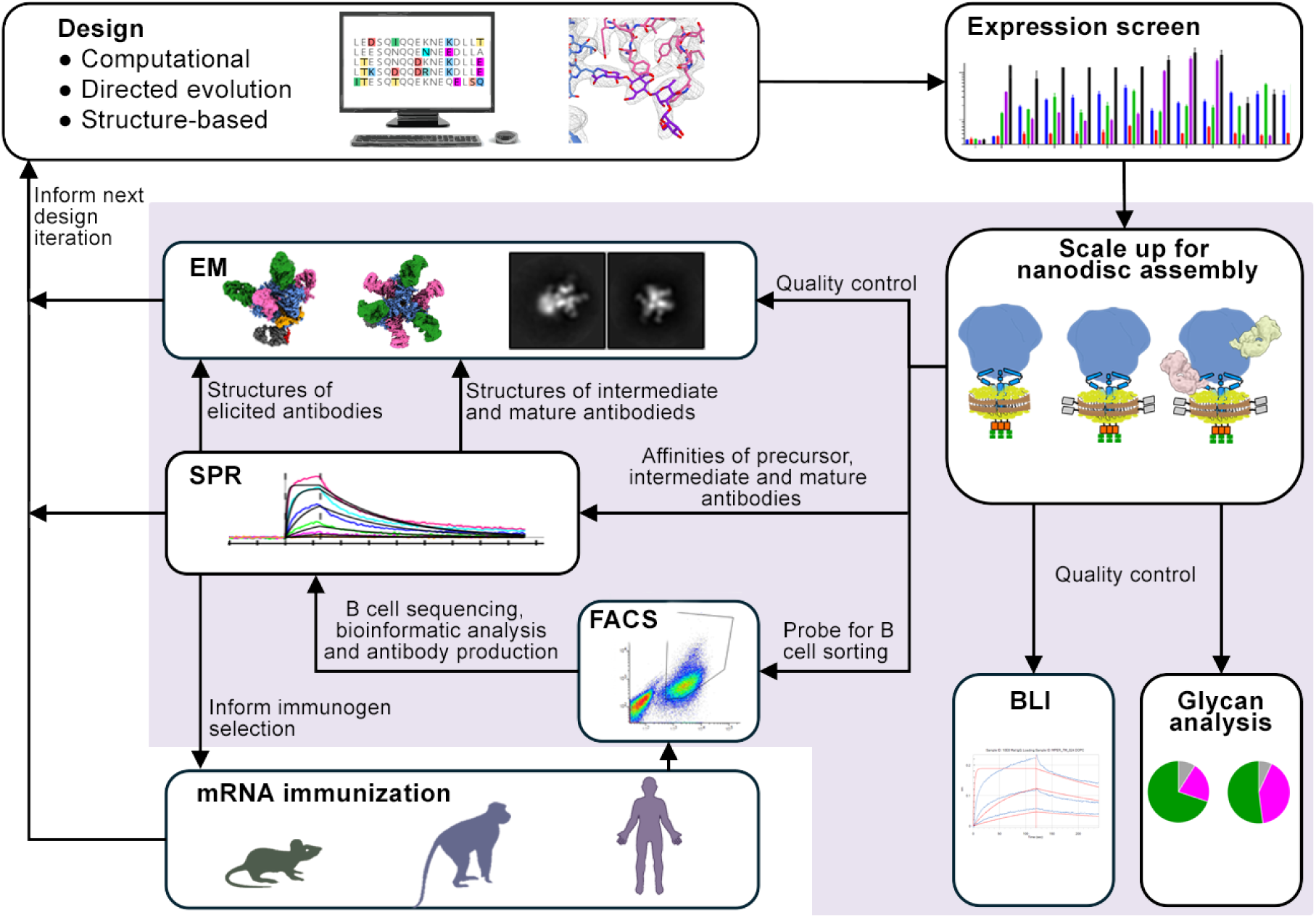
Nanodisc use in different steps in iterative rational vaccine design. Analytical methods that can directly use nanodiscs as sample material are highlighted.

**Figure 2.**
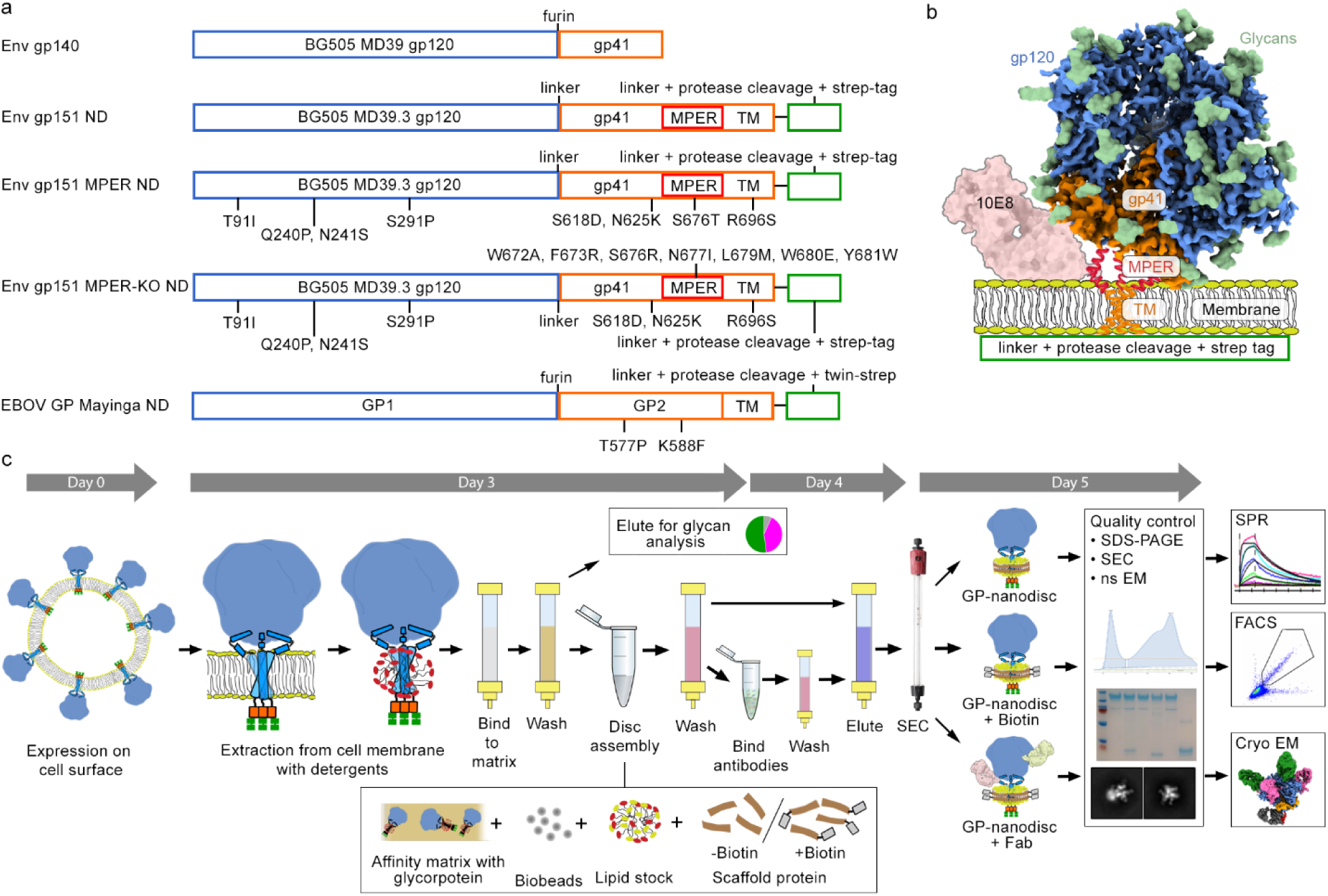
HIV Env constructs used for method development and general overview of the nanodisc assembly workflow. a) Naming and gene organization of the glycoprotein constructs used for nanodisc assembly. Introduced mutations and intracellular elements are indicated. ND refers to constructs made for nanodisc platform. b) Schematic overview of HIV Env constructs and MPER targeting antibody 10E8 binding. c) Key steps of the 5-day workflow for glycoprotein nanodisc assembly. Glycoprotein was extracted with detergent from cell surface and bound to affinity matrix. Disc assembly was performed “in-batch” while bound to affinity purification matrix. Assembled discs were then eluted for final SEC polishing purification, followed by quality control steps before subjecting to final analytical methods (SPR, FACS, cryo-EM).

**Figure 3.**
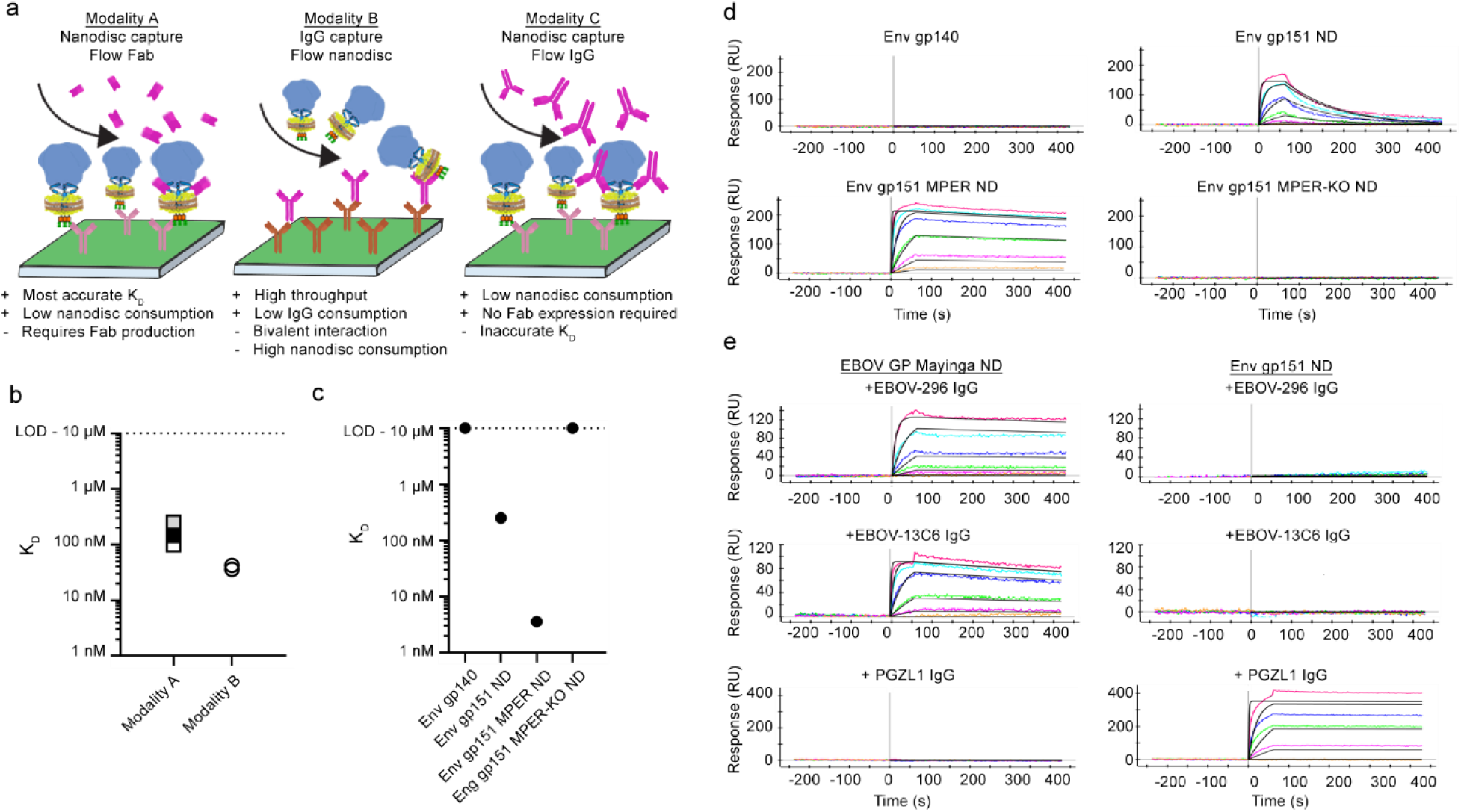
Different SPR modalities and kinetic analyses of GP nanodiscs. a) Schematic illustrations of three SPR modalities used throughout the study and their strengths and weaknesses. b) Affinity of MPER targeting 10E8 antibody to Env gp151 ND as measured with modalities A and B. Affinities with modality A were measured after storing the sample either 1 week (white), 1 month (grey) or 3 months (black) at 4°C. c) Affinity of 10E8 against Env constructs using modality A. d) SPR data presented in (c), showing increase in affinity originating from reduced off-rate in Env gp151 MPER ND. e) Modality C was used to scout Ebola specific EBOV-296 and EBOV-13C6 IgG epitope accessibility in EBOV GP nanodiscs. Data shows binding of two anti -Ebola antibodies targeting the glycan cap (EBOV-296 and 13C6). Env gp151 ND and anti-HIV MPER antibody PGZL1 were used as a negative controls.

The workflow was then piloted in an immunogen development project aiming to increase the affinity of MPER targeting antibody 10E8 and therefore guiding immune responses to MPER domain. In the native membrane context, the C-terminal MPER helix that is targeted by the heavy chain complementarity determining region 3 (CDRH3) of 10E8 is sterically occluded (Fig 2b) and one of the design goals was to improve the MPER access. To this end, N-linked glycosylation sites at positions N88, N618, and N625 (HxB2 numbering) adjacent to the epitope were removed at the base of the trimer. An additional R696S mutation that improved 10E8 binding while maintaining trimeric Env conformation was discovered by mammalian surface display library screening. Earlier structural analysis indicated that R696 resides at the intersection of the three transmembrane helices^29, 38^; therefore, this mutation was predicted to disrupt helix interactions and further open the trimer base. The resulting construct will be referred to as Env gp151 MPER ND. A negative control for MPER antibody binding and a KO FACS probe for epitope-specific B cell sorting, designated as Env gp151 MPER-KO ND, was also generated (Fig. 2a). This construct included MPER mutations W672A, F673R, T676R, N677I, L679M, W680E, Y681W, and W683D, which were previously reported to collectively prevent the binding of MPER-targeting antibodies while preserving the overall trimeric structure of Env and other bnAb epitopes^24^, or screened for this study to improve the expression construct. Prototypical Ebola GP immunogen EBOV GP Mayinga ND was based on a construct with trimer stabilizing mutations T577P and K588F^39^, to which native EBOV GP TM domain and the same intracellular elements as in the HIV Env constructs were added. Assembled nanodiscs generally exhibited broad peaks in SEC spectra indicative of extensive glycosylation and variation in hydrodynamic radius that may result from positional heterogeneity of Env in the disc, and >95% purity and bands of the expected molecular weight as examined by SDS-PAGE (Supplementary Fig. 1a-b). Ns EM 2D class averages revealed features typical of nanodiscs with trimeric GP with or without Fabs (Supplementary Fig. 4a). In addition to the MPER targeting immunogen constructs and EBOV GP Mayinga ND, a germline-targeting HIV vaccine candidate N332-GT5 gp151^40, 41^ was assembled into nanodiscs using the standardized workflow to test general applicability (Supplementary Fig. 2a). Mass spectrometric glycan analysis using the DeGlyPHER method^42^ was done prior to and after nanodisc assembly to ascertain site-specific glycan occupancy and processivity of Env constructs. This analysis confirmed the presence of all expected glycans as well as a high proportion of complex glycans in transmembrane constructs characteristic of native, transmembrane HIV Env (Supplementary Fig. 1c). Germline-targeting immunogen N332-GT5 showed a higher proportion of complex type glycans in soluble version of the immunogen, matching closely the glycan composition of the transmembrane version. All transmembrane Envs showed higher glycan occupancy in gp41 subunit.

### GP nanodisc binding kinetics to antibodies measured by SPR

SPR is an essential method in iterative vaccine design that measures antibody affinities to engineered immunogens, assessing the effect of the designed features as well as affinities of antibodies induced by vaccination (Fig 1). We set to establish scalable and reproducible conditions for measuring antibody binding kinetics against transmembrane GPs in nanodiscs in three distinct SPR modalities (Fig. 3a). Modality A utilized an anti-affinity tag capture strategy in which anti-strep-tag antibody was covalently immobilized on the chip surface, glycoprotein nanodiscs were captured using the intracellular strep tag, and Fab analytes were employed to study monovalent interactions. Modality B employed a low surface density IgG capture strategy, in which anti-human IgG capture antibody was first covalently immobilized, and glycoprotein-specific IgG ligands were captured at reduced capture time to limit the ligand density. Glycoprotein nanodiscs were then injected as analytes. In modality C, nanodisc capture is followed by IgG flow as analyte instead of Fabs, allowing a high throughput scouting approach with low nanodisc sample consumption but with reduced kD accuracy due to increased bivalent interaction. Establishing these three modalities for nanodisc GPs allowed flexibility in experiment design based on the need in iterative vaccine development step and sample availability.

Next, we used the GP nanodisc SPR modalities to establish baseline kinetics for neutralizing and non-neutralizing monoclonal HIV Env antibodies spanning diverse epitopes (Fig. 3b-d, Supplementary Fig. 2b-d). Modality A measured 250 – 100 nM affinity for 10E8 to Env gp151 ND design base construct (Fig. 3b) and slightly higher affinity of 36 – 42 nM using modality B, indicating that the low-density capture reduced but did not eliminate the avidity effects of Modality B. Using modality A, we observed 70-fold increase in 10E8 affinity (250 nM vs 3.6 nM) to Env gp151 MPER ND as a result of removal of the MPER epitope-proximal glycans and the mutation R696S in TM (Fig. 3c). Kinetic analysis indicated that the improved affinity in the engineered immunogen was primarily attributed to a decreased off-rate (8.1 × 10^−3^ s^−1^ for Env gp151 ND vs 3.5 × 10^−4^ s^−1^ for Env gp151 MPER ND. Fig. 3d). Incorporation of MPER-KO mutations completely abrogated 10E8 binding. Additional MPER targeting bnAbs and bnAbs targeting other epitopes on the surface of Env were also tested for more comprehensive antigenic profiling. Expectedly, this confirmed binding of MPER targeting antibodies to Env gp151 ND and no binding to Env gp140 (Supplementary Fig. 2b), and similar affinities of non-MPER bNabs to soluble Env gp140 and Env gp151 ND (Supplementary Fig. 2c), with reduced or absent binding of non-neutralizing antibodies RM19R and RM20A3 to Env gp151 ND as compared to Env gp140 (Supplementary Fig 2d). These two antibodies target the base of the trimer, which is exposed in Env gp140, whereas in Env gp151 ND access is presumably restricted by MPER and TM domains as well as the lipid surface of the nanodisc. Finally, we employed modality C to confirm the binding of two Ebola GP glycan cap specific antibodies EBOV-296 and 13C6 to EBOV GP nanodiscs (Fig. 3e). Collectively, these data demonstrate that GP nanodiscs and SPR can be used to characterize the antigenic landscape of the designed transmembrane immunogen, inform immunogen selection for in vivo studies, identify suitable complexes for structural studies by cryo-EM and support GP nanodisc use as B cell sorting probes in FACS (Fig 1).

### FACS sorting of B Cells from immunized animal models using nanodisc GP probes

Single B cell sorting and sequencing is routinely used to analyze B cell responses induced in vivo. Typically, biotin-tagged variants of the immunogen are used with streptavidin. This allows conjugation with multiple fluorochromes to increase selection specificity. By including a matching sorting probe with KO mutations in the targeted epitope, epitope-specific responses can be separated from off-target responses against unrelated epitopes. Soluble GP protein constructs solely consisting of the ectodomain are most often used as antigenic baits. For immunization studies where membrane-proximal epitopes are of interest, such as the HIV MPER, soluble probes would not be able to extract all appropriate responses to the targeted epitope due to several challenges which include 1) the hydrophobicity of the epitope regions which make bait production challenging, 2) non-native or incomplete structural conformation of the epitope in shorter peptide representations of the antigen, 3) missing lipid membrane components which constitute a portion of the native epitope. Here, we used biotinylated MSP1D1 scaffold to generate nanodisc GP FACS probes. We first tested Env gp151 ND probe binding to FACS compensation beads conjugated to 10E8, DH511 or BG18 bnAbs (Fig. 4a). Background signal was measured as binding to either MSP1D1 scaffold protein alone, empty nanodisc assembled with DOPC lipids, or to empty nanodisc with mixture of neutral and charged lipids. Env gp151 ND exhibited substantially higher signal compared to controls, confirming that adequate separation from background signal can be detected. We detected no binding with any of the tested probes to unimmunized C57BL/6 (B6) mouse splenocytes, and a strong signal against engineered Ramos cells expressing the bnAb VRC01 as its B cell receptor (BCR) ^43^. This data indicated that Env nanodisc probes were suitable for detection of B cells from mice by flow cytometry that specifically target Env.

**Figure 4.**
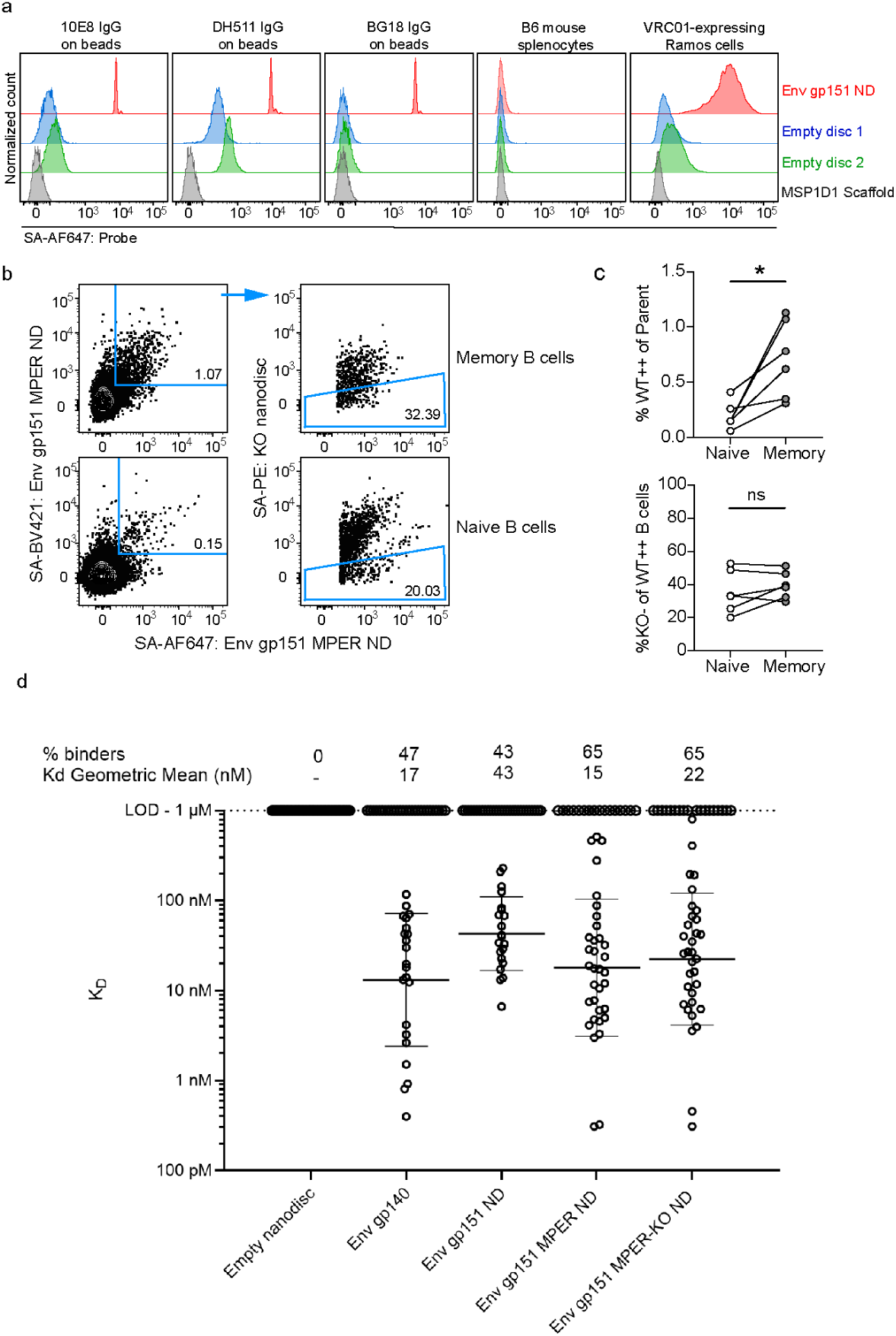
HIV Env nanodisc FACS probe validation and pilot use in pre-clinical mouse model. a) Nanodiscs were tested for binding to HIV MPER bnAbs 10E8 and DH511 and gp120-specific bnAb BG18 coupled to FACS compensation beads, B6 mouse splenocytes were used as negative control cells, and VRC01 expressing Ramos cells as positive control cells. Empty nanodiscs were assembled with either DOPC lipids (blue) or a mixture of neutral and charged lipids (green). b) Memory (CD19^+^IgM^−^IgD^−^) and naïve (CD19^+^IgM^+^IgD^+^) B cells from mice 6 weeks after immunization with mRNA-LNPs encoding Env gp151 MPER ND nanodisc were analyzed by flow cytometry. Antigen-specific (Env gp151 MPER ND^++^) and epitope-specific (KO^−^) B cells were detected using immunogen-matched WT and 10E8-epitope KO nanodisc probes and WT nanodisc^++^ cells were sorted for BCR sequencing. The complete gating strategy is shown in supplementary Fig. 3. c) Percent antigen-specific memory B cells in the naïve vs. memory compartments of each immunized mouse (top), and proportion of 10E8 epitope-specific (KO^−^) cells within each antigen-specific population. Paired t-test, *p < 0.05, ns: p> 0.05. d) Sorted cells were sequenced and selected antibodies were purified for affinity measurement by SPR using modality B.

We then proceeded with a pilot immunization experiment in mice to explore if vaccine elicited antigen-specific B cells could be identified using Env nanodiscs probes. We utilized transgenic mice expressing the human D3-3 and J_H_6 genes that allow mice to express long HCDR3s, including HCDR3 precursors of HIV MPER bnAbs 10E8, DH511 and LN01^24^. Mice were intramuscularly injected with 10 µg of mRNA-LNPs encoding HIV Env gp151 MPER ND construct. Animals were sacrificed, and spleens and lymph nodes were harvested 6 weeks post immunization. B cells were stained with immunogen-matched (WT) nanodisc probes on two distinct fluorophores and Env gp151 MPER-KO ND nanodisc probe with KO mutations in the MPER to identify MPER-specific cells (Fig. 4b and Supplementary Fig. 3). Among class-switched memory B cells (defined as CD19^+^IgD^−^IgM^−^), 0.7% of cells were determined to be antigen-specific, compared to 0.15% within the IgD^+^IgM^+^ naïve B cell population (Fig. 4c). This ∼4.7-fold increase was statistically significant (p=0.0177), indicating that the mRNA immunization elicited HIV Env gp151 MPER -specific responses that could be detected by Env nanodisc probes. The proportion of WT Env nanodisc-binding B cells that selectively bound to the 10E8 epitope (%KO^−^ of WT^++^ B cells) was indistinguishable between the naïve and memory B cell compartments, suggesting that the immunogen did not elicit MPER-specific responses, and that further engineering or inclusion of a prior priming immunogen in the immunization regimen would be required to achieve that capacity.

Antigen-specific (WT^++^) mouse memory B cells were sorted and their BCRs sequenced to assess the robustness of nanodisc probes in the selection of antigen-specific BCRs. Eight antibodies from each animal were randomly selected for monoclonal antibody production. An additional 12 antibodies with long (>=19 AA) CDRH3s were selected and duplicate antibodies (containing identical heavy- and light chains) were removed, resulting in 54 total antibodies. Genes encoding these antibodies were synthesized and 49 out of 54 antibodies were successfully expressed, purified and used in SPR modality B to measure affinities against Env nanodiscs (Fig. 4d). At 1 μM analyte concentration, no non-specific responses to empty nanodiscs were detected. Out of the tested antibodies, 47% bound to soluble gp140 matching the immunogen ectodomain and 65% to Env gp151 MPER ND or Env gp151 MPER-KO ND. Considerable binding to ectodomain, similar binding to MPER targeting immunogen and its MPER KO version and overall similar geometric mean affinity to tested Env nanodiscs provided evidence that the elicited antibodies were not targeting the MPER peptide but bound to the exposed base epitope or other off-target epitopes. Nevertheless, using the nanodiscs as FACS sorting probes and in SPR, we were able to sort and measure affinities of several antibodies that were specific to the transmembrane Env gp151 version (e.g. Ab_38, Ab_48, Ab_79. Supplementary Fig. 3b).

Non-human primates (NHPs) are an important model for late-stage pre-clinical vaccine development and are often used as final validation before clinical trials. Rhesus macaques, like humans, have extensive genetic diversity within the immunoglobulin loci and are also capable of making long CDRH3s ^44–46^. To test whether nanodisc-based FACS tetramer probes could be successfully used in RMs, experiments were carried out using PBMCs from animals immunized previously with soluble Env MD39^47^. PBMCs were stained with the nanodisc probes to determine whether there was substantial background binding to the lipid component of the nanodisc probes. Pre-immunization PBMCs showed low background binding of IgD^−^ memory B cells to nanodisc, while PBMCs from 2-weeks post an MD39 booster immunization showed high binding to the nanodisc probes (Fig. 5a). Next, the nanodisc probes were compared directly to soluble Env gp140 protein probes in RMs that had been immunized with MD39.3 gp151 mRNA (matching Env gp151 ND sequence) 4-weeks prior^48^. Across two animals there were no differences in the frequency of tetramer^++^ cells between the Env gp151 ND nanodisc and Env gp140 soluble protein probes (Fig. 5b). Taken together these data show that nanodiscs bearing stabilized HIV Env constructs can be deployed as tetramer probes for the evaluation of pre-clinical HIV B cell responses in NHPs.

**Figure 5.**
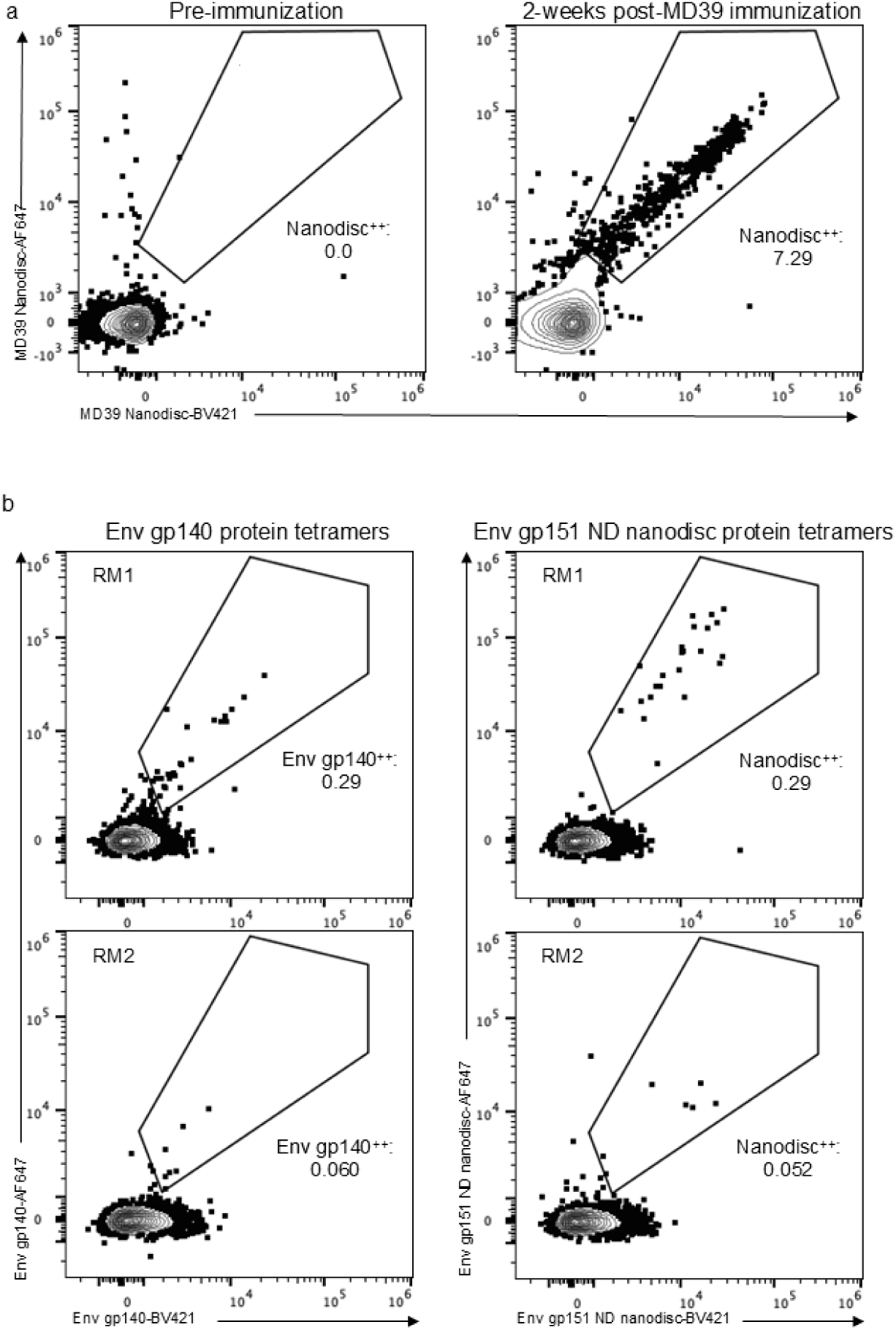
HIV Env nanodisc probe validation for sorting NHP cells. a) Flow cytometry plots showing CD20^+^IgD^−^ memory B cells stained with Env gp151 ND nanodisc on two different fluorophores. Pre-immunization and 2-weeks post-immunization PBMCs from the same RM are shown and the frequency of Nanodisc^++^ cells of IgD^−^ cells is listed. b) Flow cytometry plots showing CD20^+^IgG^+^ memory B cells stained with either soluble Env gp140 protein or Env gp151 ND nanodisc on two different fluorophores. PBMCs from two different RMs 4-weeks post mRNA Env gp151 immunization were split and stained with Env gp140 protein tetramers (left column) or matching nanodisc Env gp151 ND tetramers (right column). The frequency of tetramer^++^ cells of IgG^+^ cells is listed for each plot.

### GP nanodiscs in cryo-EM structural studies

To demonstrate how structural studies of nanodisc GPs can inform vaccine design, we set out to solve the structure of prototype MPER vaccine candidate Env gp151 MPER ND in complex with 10E8 antibody by cryo-EM. We included two other HIV bnAbs: VRC01, which recognize the CD4 binding site, and BG18, which targets the N332 glycan supersite. This antibody cocktail complex serves three purposes: First, the added Fabs improve the orientation distribution of particles on cryo-EM grids. Second, they will stabilize the trimer and prevent protomers from dissociating. Lastly, a functional HIV vaccine will likely require elicitation of bnAbs targeting several sites to achieve sterilizing immunity. A trimer boost would therefore benefit from being able to engage three or more sites (i.e. N332, CD4 and MPER) in a single immunogen. The high-resolution structure of the 10E8 epitope has thus far been resolved only with x-ray crystallography and in complex with MPER peptide alone^49–52^. From our cryo-EM data we produced two reconstructions: the complex with two 10E8 Fabs at 4.3 Å, and the complex with a single 10E8 Fab at 3.5 Å resolution, which enable atomic level interpretation. We noted that while 10E8, VRC01 and BG18 were all added at the same step after the nanodisc batch assembly at ∼10x molar excess, all VRC01 and BG18 sites were occupied but 10E8 binding site was partially underoccupied in all reconstructions, suggesting steric hinderance of the MPER epitope with increasing 10E8 occupancy (Fig. 6a-c). In all reconstructions, 10E8 is wedged between the Env ectodomain and nanodisc lipid bilayer. MPER antibody induced Env tilt angle was evaluated by aligning the low pass filtered maps to the nanodisc representing the bilayer plane. A single 10E8 Fab appears to be accommodated with minimal Env tilting while binding of a second 10E8 Fab pushes Env into a more tilted position (Fig. 6b). Unlike in our previous WT Env AMC011 nanodisc cryo-EM reconstruction, density for the TM domain remained undetected, presumably due to the R696S mutation disrupting the stabilizing connection point of the three helices in the middle of the bilayer^29^. The single 10E8 class was subjected to 3D variability analysis (3DVA), which revealed distinct conformations of the MPER helix with ∼30° lateral twist of the Fab in conjunction with an apparent dislocation of the adjacent protomer (Supplementary Fig. 4c-d. Supplementary video 1).

**Figure 6.**
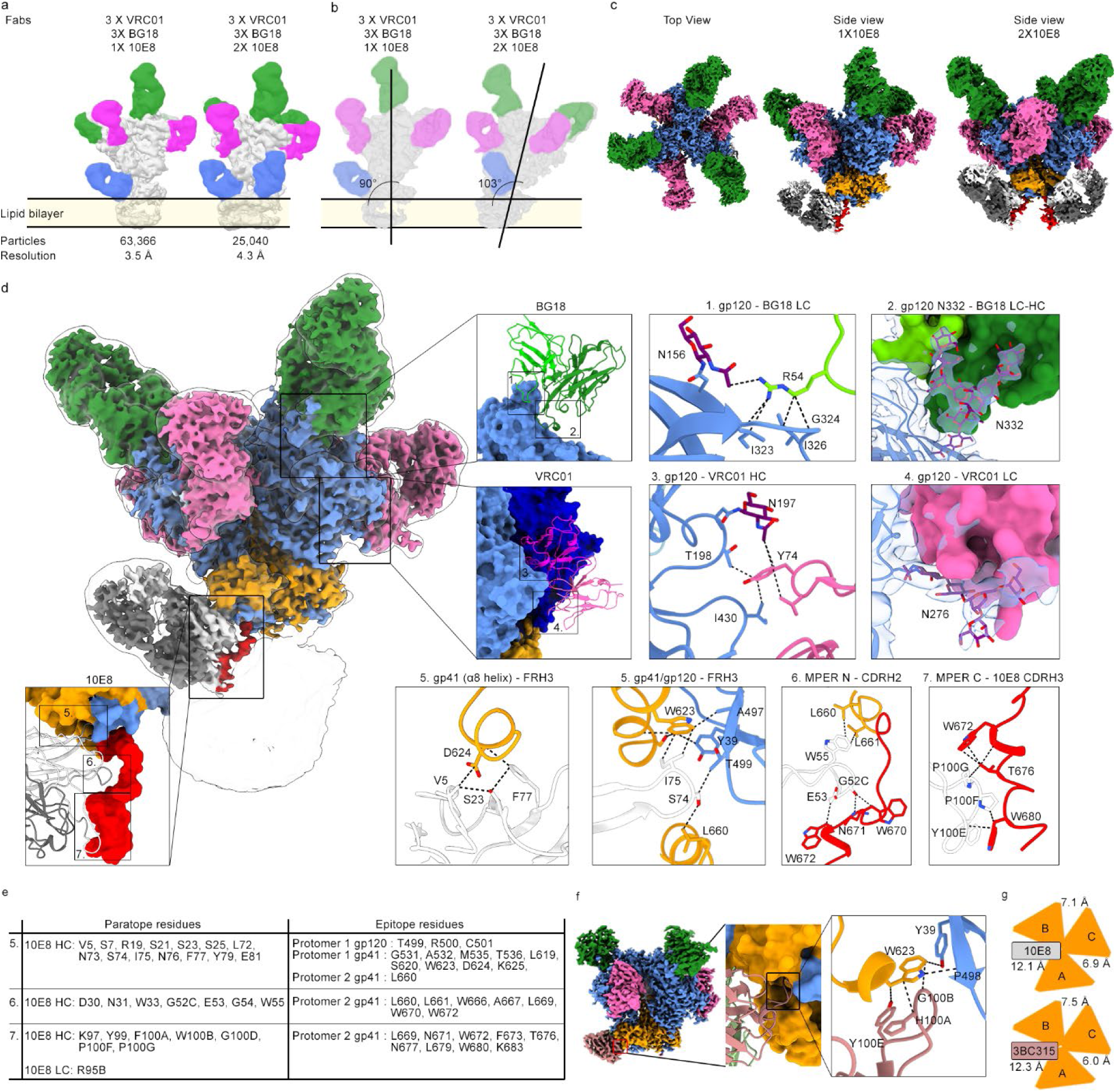
Structure of Env gp151 MPER ND in complex with bnAbs BG18, VRC01 and 10E8 resolved by cryo-EM. a) Low pass filtered maps of different Fab occupancy states and the location of the bilayer. BG18 Fab is highlighted in green, VRC01 in pink and 10E8 in blue. b) 90° clockwise rotated side views showing the Env tilt angle in relation to lipid bilayer surface. c) Top view and side views of highest resolution maps of both 10E8 occupancy states. Densities are highlighted as follows: gp120 in blue, gp41 in orange, BG18 in green, VRC01 in pink, HC of 10E8 in white, LC of 10E8 in dark grey and MPER peptide in red. d) Highest resolution reconstruction with single 10E8 was used for model building. Insets show interfaces of the three bound bnAbs and key amino acid and glycan interactions with BG18 HC (dark green) and LC (light green) in insets 1-2, VRC01 HC (magenta) and LC (pink) in insets 3-4, and 10E8 HC (white) and LC (dark grey) in insets 5-7. Glycans are highlighted in purple. Insets 2 and 4 exemplify the electron density used to built interacting glycans (transparent blue. e) Epitope-paratope analysis of 10E8 binding interface shown in d. Contacting residues are separated into three components corresponding insets 5-7. f) Binding pocket of 3BC315 (salmon) CDRH3 with key interactions highlighted. g) gp41 protomer distance analysis in the nanodisc complex structure with 10E8 Fab, and soluble Env structure with 3BC315 showing widened interface between protomers A and B due to antibody binding.

Using the model of Env gp151 MPER ND complex, we were able to resolve novel structural details of the 10E8 epitope including a binding pocket for 10E8 antibody framework region 3 (FRH3), formed by Env gp120 and gp41 subunits (Fig. 6d, inset 5). We identified 14 AA in 10E8 contacting the C -terminus of gp120, helices α6 – α8 of gp41 in protomer 1, and L660 of gp41 in protomer 2. These contacts are either in the binding pocket or in the helices forming a collar around the termini of gp120. Strikingly, this binding pocket appeared to be targeted by the CDRH3 of another HIV bnAb 3BC315, that was shown to accelerate trimer dissociation^53^. To confirm this similarity in engaging the binding pocket, we solved the cryo-EM structure of Env gp140 (BG505 MD39.3) in complex with BG18, VRC01 and 3BC315 Fabs to 3.1 Å resolution (Fig 6f, Supplemental Fig. 4f-g). 3BC315 engages the binding pocket mainly with H100A, G100B and Y100E at the tip of the CDRH3. In 10E8, the interaction is driven by I75, S74 and N76 of FRH3 (Fig 6d, inset 5). The observed similarity of the binding pocket engagement led us to propose that these antibodies may share a common mechanism for trimer destabilization. As part of the binding pocket, both antibodies contact W623 in gp41 (Fig 6d, inset 5 and 6f), which is >99% conserved across >5,000 diverse HIV strains from the LANL database (http://www.hiv.lanl.gov), and has previously been shown to be part of a tryptophan clasp contributing to stabilizing four-helix collar around the termini of gp120^54^. Engaging the binding pocket leads to widening of the gp41 protomer-protomer gap (Fig. 6g), which would eventually lead to destabilization of the contacts in the collar around the termini of gp120 and premature triggering of the fusion peptide. This would normally occur during virus entry associated Env rearrangements that follow CD4 receptor binding. Another novel structural feature of the 10E8 antibody was the dynamic remodeling of the N-terminal MPER helix, most likely driven by interactions with CDRH1 (D30, N31 and W33) and CDRH2 (G52C, G53, G54, W55. Fig. 6d-e. Supplementary video 1). Root mean square deviation (RMSD) of this CDRH1 and CDRH2 10E8 paratope region is 1.52 Å compared to a published x-ray structure of 10E8 Fab in complex with a gp41 peptide (PDB 4G6F), indicating that epitope presentation in a more native context recapitulates interactions more faithfully than epitope peptide alone. In contrast, the RMSD of FRH3 between the nanodisc cryo-EM complex and peptide-bound crystal structure is 0.46 Å. While the 10E8 crystal structure fit well into the cryo-EM map (Supplementary Fig. 4e), we calculated an RMSD of 9.59 Å of the entire MPER domain between the two structures, from the N-terminal portion of the MPER that is connected to the rest of the ectodomain in the nanodisc Env cryo-EM structure. Taken together, the comparison to previous crystal structures of 10E8 shows that native membrane environment of Env is required for complete structural characterization of the MPER targeting antibody. Observations of the native, dynamic binding mode of an MPER antibody, and structural information of the entire side chain contact network will inform the development of next-generation MPER targeting vaccines.

## Discussion

Here we present a rigorously optimized glycoprotein nanodisc platform for vaccine design. We show that the assembled GP nanodiscs enable characterization of transmembrane glycoprotein vaccine candidates with key methods in rational, iterative vaccine development that have traditionally been available only to soluble mimics. Applicability of the platform is demonstrated by using SPR to measure antibody kinetics, FACS to characterize new vaccine designs and in vivo antibody responses from preclinical experiments, and cryo-EM to provide structural guidance for immunogen design.

In addition to confirming the assembly of EBOV GP in nanodiscs with two Ebola-specific antibodies, we characterized the kinetic landscape of several HIV antibodies binding to HIV Env in nanodiscs. These confirmed that glycan shield modifications and transmembrane domain point mutation identified through directed evolution increased MPER-targeting bnAb 10E8 affinity to prototypical HIV Env MPER-targeting immunogen Env gp151 MPER ND as compared to its design predecessor. While the in vivo responses to the engineered immunogen in humanized mouse model appeared to be outside the targeted C-terminal region of MPER, the identified antibodies will provide guidance for further immunogen engineering. Potential targets of the extracted antibodies include the engineered glycan hole or off-target epitopes at the trimer base. Base of the trimer is partly occluded by the membrane, but some off-target epitopes may still be accessible. This was also indicated by EM-based polyclonal epitope mapping (EMPEM) showing that transmembrane gp151 Env administered as mRNA still elicits base-binding antibodies^15,48^. Importantly, using the biotinylated GP nanodisc as FACS probe, we were able to extract antibodies targeting sites that are not fully present in the soluble form of the immunogen. While NHP immunization was beyond the study’s scope, we also confirmed GP nanodiscs can be used to identify antigen-specific responses from NHPs, facilitating future studies. Two promising approaches for overcoming the challenges in eliciting MPER targeting antibodies have been recently described, namely priming with germline-targeting immunogen 10E8-GT12^24, 55^ or immunizing with MPER peptide alone in liposome formulation^25^. Both solutions present a more accessible MPER peptide for antibody CDRH3. These immunogens can be followed by a next generation Env boost immunogen designs presenting the complete epitope to guide the maturing antibody toward breadth and higher neutralization potency. The analysis and nanodisc GP FACS methods presented here provide vital guidance for these designs.

The platform enabled us to determine the structure of Env gp151 MPER ND to 3.5 Å, leading to discovery of structural features that will inform the next generation of immunogens. The structure confirms three HIV bnAb binding sites targeted in ongoing vaccine development projects can be engaged by the same immunogen^56,24, 44^. This further demonstrated that the nanodisc platform has the potential to be used in cryoEMPEM to resolve structures of membrane proximal antibody epitopes from a mixture of specificities from post-immunization serum^57^. As multiple epitopes are now being engineered into Env trimers for germline targeting, it is critical to understand if and how they may compete or synergize. Additional value in using GP nanodiscs is that, in contrast to previous structural studies using soluble mimetics of Env^40, 58^, this nanodisc-based structure resembles more closely the native transmembrane protein immunogen^48^, which may be important i.e. when assessing glycan engagement in the epitope. Glycan analysis of Env Env gp151 ND and Env gp151 MPER ND agreed with earlier studies showing that when Env is expressed as transmembrane protein, glycan processing resembles more closely that of the virus particle with higher proportion of complex type glycans as in soluble forms^26, 27^. The most significant value of the Env gp151 MPER ND complex cryo-EM structure however is in characterizing the MPER epitope in unprecedented detail. We were able to resolve contacts of 10E8 bnAb with the Env ectodomain that have thus far remained elusive in structural studies. A binding pocket for 10E8 FRH3 in the gp120-gp41 interface of the ectodomain, and CDRH1, 2 and 3 contacts to the entire MPER domain ending at the lipid bilayer interface, form a continuous network of side chain interactions together spanning 29 AA in the paratope and 26 AA in the epitope. These details can now be used as a structural template to design immunogens for improved engagement of MPER antibodies. Finally, 3D variability analysis of the cryo-EM data illustrates significant remodeling of the N-terminal MPER helix induced by CDRH1 and CDRH2 of 10E8. While care should be taken when interpreting flexible regions, this captures detailed snapshots of the native flexibility of the glycoprotein and MPER domain. The data depicts the range of flexibility and may offer clues how to accommodate native glycoprotein flexibility in vaccine design.

In conclusion, the GP nanodisc platform offers a scalable and reproducible solution that provides a more complete and native-like environment for transmembrane glycoprotein vaccine designs. This platform can therefore accelerate the development of next-generation vaccines and may be applicable beyond vaccine development. The described methods present significant additions to the rapidly increasing nanodisc applications in drug discovery technologies^59^. They may be used in workflows for small molecule screening, antibody discovery, and functional studies involving interactions with both extracellular and intracellular binders of any transmembrane protein.

## Methods

This method description aims to provide a complete platform that enables routine use of nanodisc approach for iterative vaccine design using viral glycoproteins. While retaining the versatility of a modular approach, each step is simplified and validated for maximizing reproducibility and scalability.

### Protein expression and nanodisc assembly

All glycoprotein and antibody constructs were codon optimized for expression in human cells and synthesized and cloned into expression vectors pHLSec (glycoproteins) or its variants pCW-CHIg-hG1, pFabCW pCW-CLig-hL2 or pCW-CLig-hk (IgG heavy chains, Fab heavy chains, lambda light chains and kappa light chains, respectively) by GenScript Biotech. Antibodies were expressed and purified from HEK293F or ExpiCHO cells according to manufacturer’s instruction using rProtein A Sepharose Fast Flow resin (Cytiva) as described earlier^24, 44^.

Soluble Env GP were purified as described earlier^40^. Transmembrane glycoprotein purification method was modified from the earlier protocol^29^. For one liter of transfected cells, 1 mg of DNA was mixed with 25 ml of OPTI-MEM medium and sterile filtrated through 0.22 µm filter unit. 3 mg of PEI MAX (Polysciences, #24765) was separately mixed with 25 ml of OPTI-MEM, filtrated and then combined with the DNA. Combined DNA and PEI MAX were then added to HEK293F cells at 1 million/ml density and cells were incubated at 37 °C, 8% CO_2_, and 80% humidity under shaking at 180 rpm for 3 days. Cells were harvested by centrifugation at 1500 rcf in 4 °C for 15 min, washed with ∼300 ml cold PBS, and centrifuged again at 1500 rcf in 4 °C for 15 min. Pelleted cells were lysed with ∼5 ml of lysis buffer per gram of cells (50 mM Tris-HCl (pH 7.4), 300 mM NaCl, 0.5% TX-100). Prior to use, approximately 1 protease inhibitor cocktail tablet per 50 ml of lysis buffer, and 2 ml of BioLock (IBA Lifesciences, # 2-0205-050) per liter of cells was added to lysis buffer. Cells were lysed for 1 h at 4 °C in an overhead rotating mixer. Lysed cells were centrifuged for 1 h at 25,000 rcf, followed by filtration of the supernatant through 0.22 µm bottle top filtration unit. This resulted in cleared lysate that was passed through Strep-tactin XT 4Flow matrix (IBA Lifesciences, #2-5010-025) in a gravity flow column. Approx. 400 µL of drained matrix was used per liter of cells. Matrix was then washed three times with 2 ml of wash buffer 1 (50 mM Tris-HCl (pH 7.4), 300 mM NaCl, 0.1% (w/v) CHAPS, 0.03 mg/mL deoxycholate), three times with 2 ml of wash buffer 2 (50 mM Tris/HCl pH 7.4, 500 mM NaCl, 1 mM EDTA, 0.1% DDM, 0.01% CHS, 0.03 mg/ml deoxycholate) and three times 2 ml of wash buffer 3 (50 mM Tris/HCl, pH 7.4, 150 mM NaCl, 0.02% DDM, 0.002% CHS, 0.03 mg/ml deoxycholate). After this, matrix with detergent solubilized GP was transferred to a 5 ml test tube for nanodisc assembly step.

Lipid stocks were prepared prior to disc assembly in 1 mM final concentrations. Premade 10% DDM/ 0.1% CHS stock (Anatrace, #D310-CH210) was used throughout the study. All lipids in this study were prepared from chloroform stocks (Avanti Polar Lipids). Chloroform was first evaporated under gentle nitrogen stream until transparent film was formed around the glass vial. Lipid film was then rehydrated for 1 h in room temperature in lipid rehydration buffer (50 mM Tris/HCl, pH 7.4, 150 mM NaCl, 0.1% DDM, 0.01% CHS) in a volume bringing the lipid stock concentration to 5 mM. After thorough mixing by vortexing, this stock was diluted further to final lipid stock solution at 1 mM concentration and sonicated with microtip (20-25% intensity, 50% time cycles in 4 °C) until clear. Most lipid stock solutions cleared within 10 – 20 min but with some lipids DDM/CHS amount was increased up to 1%/0.01%. 1 mM lipid stocks were aliquoted and stored up to 6 months at -20°C. A lipid mix roughly following HIV particle lipid composition was used throughout the study and contained 45% DOPC, 7% DOPS, 7% DOPA, 7% DOPE, 7% DOPG, 5% sphingomyelin, 2% PI(4,5)P2, and 20% CHS. Scaffold protein was either produced in-house using standard *E. coli* expression method or purchased from Sigma-Aldrich (membrane scaffold protein 1D1 #M6574 or biotinylated membrane scaffold protein 1D1 BTN #MSP13).

Nanodisc batch assembly was done as follows. Strep-tactin matrix with GP bound was drained by gravity flow, after which gravity flow column was capped and 400 µl of wash buffer 3 was added to the matrix. 50% matrix slurry was then transferred to a 5 ml test tube. Lipids were added at approx. 350X molar excess and scaffold at 6X molar excess in relation to the glycoprotein. Glycoprotein amount bound to the matrix was estimated with a separate initial experiment where protein was eluted after the last wash step. Minor variations were made to glycoprotein:scaffold and glycoprotein:lipid ratios without significant difference in the final yield. Matrix with glycoprotein, lipids and scaffold was incubated for 1 h at 4°C prior to addition of SM-2 bio-beads (Bio-Rad, #1523920). Bio-beads were activated according to the manufacturer’s instructions in methanol followed by extensive wash with deionized water. Bio-beads were added to approx. 50% v/v with the nanodisc assembly mixture. i.e. if the total volume of the assembly reaction was 1.5 ml, final volume after bio-bead addition in the 5 ml test tube was ∼3 ml. Detergent was then removed by incubation in a rotating shaker overnight at 4°C. The following day an additional ∼20% of the total reaction volume of fresh bio-beads were added, followed by an additional 1 h incubation at room temperature to ensure complete removal of detergent. Next, contents of the test tube were transferred to a new gravity flow column with a pipette tip with the first ∼2 mm cut off to allow pipetting biobeads and matrix. Mixture was washed in column twice with 4 ml of TBS and eluted with Strep-tactin elution buffer BXT in 500 µl fractions. A280 of eluted fractions were monitored and fractions containing protein were pooled and concentrated to <500 µl for SEC performed using a Superose 6 Increase 10/300 column (Cytiva) at 0.75ml/min in TBS. SDS-PAGE and ns EM analysis were used to confirm presence of glycoprotein and scaffold protein, typical disc appearance in ns EM 2D class averages and glycoprotein on disc.

For structural biology purposes, Fabs were added to the GP nanodiscs after the assembly while nanodisc was still bound to strep-tactin matrix. After the second addition of bio-beads, each Fab was added at ∼10X molar excess and the mixture was incubated for an additional 2h at room temperature. Complex elution and following steps were identical as described above for unliganded glycoprotein nanodiscs. Fractions were pooled after SEC and sample was concentrated either to 1 mg/ml for FACS, SPR and other purposes, or up to 11 mg/ml for cryo-EM sample preparation.

### Identification of R696S mutation on HIV Env by mammalian directed evolution

A 250 amino acid long site-saturation mutagenesis (NNK) scan of gp41 was synthesized at SGI-DNA. A fragment of the library containing NNKs from positions T606 to F699 (94 positions) was cloned into BG505 MD39.2 gp151. This library was stably integrated into 293T-rtTA3G cell line using the lentivirus-based mammalian display protocol described previously^34, 40,35^. After inducing library expression overnight, the cells were stained with 10E8 IgG and PGT145 Fab and those with highest binding to 10E8 that were also positive for PGT145 were sorted. After three rounds of sorting, the library DNA was extracted and sequenced by Sanger sequencing. 8 of 19 sequences had a mutation at position 696, which was mutated from R to S, A, V, I and L. The most frequent mutation, R696S, was selected for further characterization.

### Mass spectrometry

#### Proteinase K treatment and deglycosylation

HIV Env glycoprotein (when membrane bound, denatured in 6 M urea) was exchanged to water using Microcon Ultracel PL-10 centrifugal filter. Glycoprotein was reduced with 5 mM tris(2-carboxyethyl)phosphine hydrochloride (TCEP-HCl) and alkylated with 10 mM 2-Chloroacetamide in 100 mM ammonium acetate for 20 min at room temperature (RT, 24 °C). Initial protein-level deglycosylation was performed using 250 U of Endo H for 5 µg trimer, for 1 h at 37 °C. Glycoprotein was digested with 1:25 Proteinase K (PK) for 30 min at 37 °C. PK was denatured by incubating at 90 °C for 15 min, then cooled to RT. Peptides were deglycosylated again with 250 U Endo H for 1 h at 37 °C, then frozen at –80 °C and lyophilized. 100 U PNGase F was lyophilized, resuspended in 20 µl 100 mM ammonium bicarbonate prepared in H_2_^18^O, and added to the lyophilized peptides. Reactions were then incubated for 1 h at 37 °C, subsequently analyzed by LC-MS/MS.

#### LC-MS/MS

Samples were analyzed on an Q Exactive HF-X mass spectrometer. Samples were injected directly onto a 25 cm, 100 μm ID column packed with BEH 1.7 μm C18 resin. Samples were separated at a flow rate of 300 nL/min on an EASY-nLC 1200 UHPLC. Buffers A and B were 0.1% formic acid in 5% and 80% acetonitrile, respectively. The following gradient was used: 1–25% B over 160 min, an increase to 40% B over 40 min, an increase to 90% B over another 10 min and 30 min at 90% B for a total run time of 240 min. Column was re-equilibrated with solution A prior to the injection of sample. Peptides were eluted from the tip of the column and nanosprayed directly into the mass spectrometer by application of 2.8 kV at the back of the column. The mass spectrometer was operated in a data dependent mode. Full MS1 scans were collected in the Orbitrap at 120,000 resolution. The ten most abundant ions per scan were selected for HCD MS/MS at 25 NCE. Dynamic exclusion was enabled with exclusion duration of 10 s and singly charged ions were excluded.

#### Data Processing

Protein and peptide identification were done with Integrated Proteomics Pipeline (IP2). Tandem mass spectra were extracted from raw files using RawConverter ^60^ and searched with ProLuCID^61^ against a database comprising UniProt reviewed (Swiss-Prot) proteome for Homo sapiens (UP000005640), UniProt amino acid sequences for Endo H (P04067), PNGase F (Q9XBM8), and Proteinase K (P06873), amino acid sequences for the examined proteins, and a list of general protein contaminants. The search space included no cleavage-specificity. Carbamidomethylation (+57.02146 C) was considered a static modification. Deamidation in presence of H_2_^18^O (+2.988261 N), GlcNAc (+203.079373 N), oxidation (+15.994915 M) and N-terminal pyroglutamate formation (–17.026549 Q) were considered differential modifications. Data was searched with 50 ppm precursor ion tolerance and 50 ppm fragment ion tolerance. Identified proteins were filtered using DTASelect2^62^ and utilizing a target-decoy database search strategy to limit the false discovery rate to 1%, at the spectrum level^63^. A minimum of 1 peptide per protein and no tryptic end per peptide were required and precursor delta mass cut-off was fixed at 15 ppm. Statistical models for peptide mass modification (modstat) were applied. Census2^64^ label-free analysis was performed based on the precursor peak area, with a 15 ppm precursor mass tolerance and 0.1 min retention time tolerance. “Match between runs” was used to find missing peptides between runs. Data analysis using GlycoMSQuant ^42^ was implemented to automate the analysis. GlycoMSQuant summed precursor peak areas across replicates, discarded peptides without NGS, discarded misidentified peptides when N-glycan remnant-mass modifications were localized to non-NGS asparagines and corrected/fixed N-glycan mislocalization where appropriate.

#### SPR

The kinetics and binding affinities of antibody-antigen interactions were analyzed using either ProteOn XPR36 system (Bio-Rad) or Carterra LSA. TBS pH 7.4 (20 mM Tris, 150 mM NaCl) supplemented with BSA at 1 mg/ml without detergents was used as running buffer in all experiments. In ProteOn XPR36, HC30M XanTec sensor chips were utilized. In modalities B and C, anti-human IgG (Fc) antibody (GE, #BR-1008-39) was used for capturing IgG (ligand) at low densities with nanodisc GP as the analyte. In modality A, anti-Strep-tag antibody (pAb, rabbit, GenScript, #A00875) was used for capturing Strep-tagged nanodiscs GP (ligand) followed by GP specific Fab as the analyte. Approximately 6000-8000 response units (RU) of the capture antibody were covalently attached to the sensor surface using EDC-NHS chemistry. For IgG-antigen interaction studies, approximately 50 to 100 RUs of IgGs at a concentration of 0.3 μg/ml were immobilized on each flow cell. For Fab-antigen interaction studies, approximately 300 to 400 RUs of antigen at 10 μg/ml were captured on each flow cell. Analytes were introduced to the flow cell at a rate of 30 μl/min for 3 minutes, followed by a dissociation phase of 5 minutes. Regeneration was performed with 1.7% or 0.85% phosphoric acid, each with a contact time of 60-180 seconds, repeated four times per cycle. Data analysis was conducted using ProteOn Manager software (Bio-Rad), including raw sensogram processing, interspot referencing, and column double referencing. Equilibrium or kinetic fits were performed using the Langmuir model as needed.

Kinetics and affinity of antibody-antigen interactions on Carterra LSA using CMDP Sensor Chip (Carterra) for capture IgG – flow nanodisc (modality B) was done as follows. Chip surfaces were prepared according to Carterra software instructions. In a typical experiment approx. 500-700 RU of capture antibody (SouthernBiothech, # 2047-01) in 10 mM Sodium Acetate pH 4.5 was amine coupled on CMDP chip placing special care on concentration range of the amine coupling reagents. We used N-Hydroxysuccinimide (NHS) and 1-Ethyl-3-(3-dimethylaminopropyl) carbodiimidehydrochloride (EDC) from Amine Coupling Kit (GE, #BR-1000-50). Highest coupling levels of capture antibody were achieved by using 10 times diluted NHS and EDC during surface preparation runs as compared to kit instruction (10 ml of water each to give 11.5 mg/ml and 75 mg/ml respectively according to kit instructions). Thus, the concentrations of NHS and EDC were 1.15 mg/ml and 7.5 mg/ml and the activation time was reduced to 1 min. The concentrated stocks of NHS and EDC were stored frozen in -20 °C for up to 2 months without noticeable loss of activity. The capture antibody was used at concentration 25 µg/ml with 10 minutes contact time. Phosphoric Acid 1.7% was used as a regeneration solution with 60 seconds contact time and injected three times per cycle. Concentration of ligands were approximately 1 µg/ml and contact time was 5 min. Raw sensograms were analyzed using Kinetics software (Carterra) with interspot and blank double referencing, and Langmuir model. Analyte concentrations were quantified on NanoDrop 2000c spectrophotometer using absorption signal at 280 nm. Analytes were buffer exchanged into the running buffer using dialysis. In a typical run, we covered a broad range of affinities and used two referencing practices depending on the off-rate of the ligand. For fast off-rate (faster than 9 × 10^−3^ 1/s), we use automated batch referencing that includes overlay y-aline and higher analyte concentrations. For slow off-rates (9 × 10^−3^ 1/s or less), we use manual process referencing that includes serial y-align and lower analyte concentrations. After automated data analysis by Kinetics software, we performed additional filtering to remove datasets with highest response signals smaller than signals from negative controls using R-script.

### FACS

#### Validation of Env-Nanodisc proteins as fluorescent baits

Streptavidin (SA) conjugated-antigen baits were prepared by combining biotinylated Env-nanodisc or empty nanodisc with fluorescent SA at room temperature for at least 1 hr in dark at a bait:SA molar ratio of 2:1. Control breads for FACS were generated by conjugating various bnAbs to compensation beads. Mouse anti-human IgG (BD Biosciences, # 555784) was first captured on to anti-mouse IgK compensation beads (BD Biosciences, # 552843), followed by a wash step and a secondary capture of bnAbs of interest. VRC01-expressing Ramos cells ^43^, C57BL/6 splenocytes, and the prepared bnAb-conjugated beads were incubated with the nanodisc GP:SA baits at a bait concentration of 10-50 nM in the final staining volume for 30 min at 4 °C in the dark. Data were acquired on a BD FACS Melody, and analzyed in FlowJo v10 (FlowJo, LLC).

#### Mouse immunization studies

hD3-3/JH6 mice were immunized as previously described^24^. Briefly, homozygous hD3-3/JH6 mice were injected with 10 µg (50 µl total volume) of Moderna mRNA LNPs encoding Env gp151 MPER I.M. under anesthesia (5% isoflurane induction) in the left quadriceps muscle. After six weeks, mice were euthanized with compressed CO_2_ (100%) in a clear chamber to allow for visualization of respiration and subsequent death via respiratory cessation. Blood was collected from the chest cavity prior to the removal of the spleen and lymph nodes (mesenteric, inguinal, and popliteal). Tissues were placed in 3 mL FACS buffer (1 X PBS Ca/Mg^++^ free, 1 mM EDTA, 25 mM HEPES, pH 7.0, 1% heat-inactivated FBS) on ice and tissues were disassociated using the rough ends of two sandblasted microscope slides, followed by centrifugation (460 rcf for 5 minutes at 4 °C). Red blood cell lysis was performed using 1 mL of ACK buffer (Quality Biological) for 2 minutes on ice before lysis was halted by adding 14 mL FACS buffer per sample. Post lysis and centrifugation (460 xg for 5 minutes), cells were resuspended in 3 mL Bambanker freezing medium (Bulldog Bio) prior to filtration through a cotton-plugged, borosilicate pasteur pipette into a borosilicate glass test tube. 1 mL filtered-cell solution was subsequently divided into three cryovials/mouse, which were precooled in a styrofoam rack on dry ice. Cells were stored at -80 °C for 2-7 days prior to long-term storage in liquid nitrogen. All work followed IACUC guidelines associated with animal protocol number 20-0001.

#### Mouse sample preparation and B cell sorting

Mouse frozen mouse splenoctyes and lymphocytes were thawed and stained as previously described^24, 56^. Briefly, after thawing and counting, total B cells were isolated by negative selection using the EasySep Mouse Pan-B Cell Isolation Kit (StemCell, #19844) according to manufacturer’s instructions. SA-conjugated baits were prepared as described above. Wildtype antigen baits were conjugated to BV421-SA (BioLegend, #405225) and AlexaFluor 647SA (Invitrogen, #S21374), while epitope knockout (KO) baits were conjugated to hashtagged TotalSeq-C PE SA (BioLegend, #405261). Purified B cells were stained at 4 °C in dark, with appropriate baits and antibody master-mix consisting of FITC anti-CD19 (BioLegend, #152404), BV786 anti-IgM (BD Biosciences, #743328), PerCP-Cy5.5 anti-IgD (BD Biosciences, #564273), APC-Cy7 anti-F4/80 (BioLegend, #123118), APC-Cy7 anti-CD11c (BD Biosciences, #561241), APC-Cy7 anti-Ly6C (BD Biosciences, #557661), APC-H7 anti-CD8a (BD Biosciences, #560182), and APC-H7 anti-CD4 (BD Biosciences, #560181), all used at 1:100 dilution and prepared in FACS buffer (1% v/v heat inactivated FBS, 1 mM EDTA, 1 mM HEPES in 1x DPBS). The KO bait was first added to cells with the antibody master mix for 15 minutes, followed by the addition of WT baits for an additional 30 min. Each bait was used at final bait concentration of

100 nM in the staining volume or 0.5 µg per sample. During the addition of antibody master mix, a unique TotalSeq-C anti-mouse hashtag antibody (BioLegend) was added to each sample at a concentration of 2.5 µL / up to 20 million cells. Following antibody staining, 1 mL of 1:300 live/dead stain diluted in FACS buffer (Invitrogen, #L34966) was added to each sample for an additional 15 minutes at 4 °C, then washed with 10 mL of FACS buffer. Cells were resuspended in a final volume of 500 µL FACS buffer and sorted on a BD FACS Melody. Cells were bulk sorted into a 4 °C chilled PCR plate well containing 20 µL of 0.2 µm filtered FBS, and processed for BCR sequencing via the 10x Genomics Single Cell Immune Profiling according to kit protocols, with minor modifications outlined previously^65^. Pooled libraries were sequenced on an Illumina NextSeq 2000 using a 100-cycle P3 reagent kit (Illumina, #20040559).

#### BCR sequence analysis

Sequence analysis Raw sequencing data were demultiplexed, processed into assembled VDJ contigs and counts matrix files, and assigned to specific animal IDs based on TotalSeq-C antibody hashtag counts using Cell Ranger (v6.1) and scab as previously described^65^. Gene assignment, annotation, and formatting into Adaptive Immune Receptor Repertoire (AIRR) format^66^ for paired heavy and light chain antibody sequences was performed using Sequencing Analysis and Data library for Immunoinformatics Exploration (SADIE) with a previously described custom germline reference database that included all known mouse antibody genes plus the human IGHD3-3 and IGHJ6 genes that were knocked into hD3-3/JH6 mice^24^. Eight antibodies from each animal were randomly selected for in vitro analysis, except for one animal from which only two total heavy/light pairs were recovered. In addition, 12 antibodies with long HCDR3s (>19 aa) were selected for in vitro characterization.

#### NHP Flow Cytometry

Frozen NHP PBMC samples were thawed in RPMI media with 10% FBS, 1% GlutaMAX, and 1% Penicillin/Streptomycin (R10). NHP samples used in this study were from animals that were reported in previous studies ^47,48^. The recovered cells were then stained with the appropriate antibody panel. Fluorescent antigen probes were constructed by combining fluorophore-conjugated streptavidin and biotinylated nanodisc GP or soluble Env probes with an appropriate volume of PBS in small increments across 45 minutes at room temperature (RT). Thawed cells were incubated with probes for thirty minutes at 4°C. A master mix of surface staining antibodies was then added and incubated for thirty minutes at 4°C. Cells were washed and prepared for acquisition on Cytek Aurora (Cytek Biosciences). The following reagents were used in the PBMC staining panel: Alexa Fluor 647 Streptavidin (BioLegend #405237), BV421 Streptavidin (BioLegend #405225), PE Streptavidin (BioLegend #405245), PE-Cy7 Streptavidin (BioLegend #405206), BV711 Streptavidin (BioLegend #405241), BUV615 Streptavidin (BioLegend #613013), Viability eFluor780 (1:2000, Invitrogen #65-0865-14), CD3 BV510 (1:100, BD Biosciences #569488), CD14 BV510 (1:100, BioLegend #301842), CD8a BV510 (1:100, BD Biosciences #563256), CD16 BV510 (1:100, BioLegend #302048), CD20 (1:100, BUV395 BD Biosciences #563782), IgG BV605 (1:100, BD Biosciences #563246), and IgD AF488 (1:50, Southern Biotech #2030-30).

#### EM

Negative stain EM was used for quality control of nanodisc assembly and for initial assessment of antibody complexes for cryo-EM. Glycoprotein nanodisc or nanodisc with three fold molar excess of Fab was applied to 400 mesh size Cu grid at 0.04-0.06 mg/ml concentration, blotted off with filter paper and stained with 2% uranyl formate for 60 s. NsEM data was collected on a Tecnai Spirit microscope operating at 120 keV, using a Tietz 4k × 4k TemCam-F416 CMOS camera and Leginon automated image collection software^66^. Data was processed using Relion 3.0 image processing pipeline^67^.

For cryo-EM, nanodisc was complexed with 10X molar excess of VRC01, BG18 and 10E8 Fabs after batch nanodisc assembly, while bound to affinity matrix. Eluted complex was then SEC purified (Superose 6 Increase column. Cytiva), concentrated to 8 mg/ml and frozen on graphene oxide grids (GO on Quantifoils R1.2/1.3, Cu, 400 mesh, Electron Microscopy Sciences) with 30 sec wait time, blot force of 0, and blot time of 2.5-3 s. Fluorinated Fos-Choline-8 (Anatrace # F300F) was added to sample immediately prior to freezing to a final concentration of 3 mM. GO grids, high sample concentration and fluorinated Fos-Choline-8 were necessary to prevent freeze-denaturation, ensure high particle count per micrograph and prevent orientation bias. 11,871 micrographs were collected at pixel size 0.718 Å using a Thermo Fisher Scientific Glacios 2 microscope operating at 200 kV and equipped with a Thermo Fisher Scientific Falcon 4i camera using a total dose of 44.89 e-/Å2. Automated data collection was performed using the EPU software (Thermo Fisher Scientific) and images were written in the EER frame format. Micrographs were preprocessed using cryoSPARC Live^67^, including motion and CTF correction. Briefly, particles were picked with Topaz^68^ after training the neural network with a subset of particles first picked with blob picker. Picked particles were subjected to 3 rounds of 2D classification to obtain a final particle stack of 317,792 particles. These were then extracted from micrographs using a box size of 540 pix. Maps containing one or two copies of 10E8 Fab were separated using heterogenous reconstruction and 3D classification tasks. A class with 7,964 particles containing two copies of 10E8 was refined to 4.3Å resolution. A class with 111,002 particles containing single 10E8 was examined with 3D variability analysis^69^. Further classification resulted in final, cleaned particle stack containing 63,366 particles which was then subjected to Non-Uniform Refinement^70^ and used for atomic model building. Data collection and processing stats are summarized in Supplementary Table 1. Atomic model building was initiated by docking in available structures of 10E8 Fab (PDB 4G6F), BG18 Fab (PDB 6DFG), VRC01 Fab (PDB 3NGB) and an AlphaFold3^71^ -generated model of the trimer into the map using ChimeraX^72^. Manual adjustment was done using Coot^73^ and further refinement was done using Phenix Real Space Refine^74^ and Rosetta Relax^75^. Final model statistics are summarized in Supplementary Table 1.

The soluble gp140 (MD39.3) cryo-EM structure was done as follows: VRC01, BG18 and 3BC315 Fabs were added at 4.3X molar excess to purified Env glycoprotein. The complex was purified using a HiLoad Superdex 200 16/600 pg column (Cytiva) and concentrated to 10 mg/ml. Samples were vitrified using a Thermo Fisher Scientific Vitrobot Mark IV operating at 100% humidity, 4°C and 3-6 s blot times, on UltrAuFoil 1.2/1.3-300 holey gold film grids (Electron Microscopy Sciences). Prior to grid application, lauryl maltose neopentyl glycol (LMNG) was added to a final concentration of 0.005 mM. Imaging was performed using the same microscope and conditions as for Env gp151 MPER ND complex. After initial cryoSPARC particle picking using Blob Picker, templates were selected from 2D classification and used for Template Picker. A total of 886,907 particles were extracted using a box size of 576 downsampled to 144. After additional rounds of 2D classification, an Ab Initio Reconstruction was performed, followed by Non-Uniform refinement. A single 3BC315 Fab was visible in the C1 reconstruction. A mask was created over 3BC315 and focused 3D classification performed to remove trimers with no 3BC315 particles. A final stack of 243,503 were re-extracted and downsampled to box size 400 (resulting in pixel size of 1.034 Å) and subjected to Non-Uniform Refinement which resulted in a 3.1 Å reconstruction. Model building was performed as above and final model statistics are summarized in Supplementary Table 1.

Maps and models have been deposited to the EMDB and PDB, respectively, under the accession codes listed in Supplementary Table 1.

## Acknowledgements

We thank Nicole Doria-Rose for providing the VRC01 expressing Ramos cell line. We thank Brian Briney for providing access to Illumina NextSeq 2000 sequencer. We thank Daniel Murin for providing Ebola GP specific antibodies.

## Author contributions

K.R. designed nanodisc assembly workflow, performed cryo-EM imaging, data processing and structural analysis, and wrote the manuscript. A.L. and O.K. established conditions and conducted SPR assays. G.O. Supervised cryo-EM imaging and data processing and built 3D models. C.F. performed and J.H.L. and D.S. supervised establishing conditions for mouse B cell FACS experiments. J.M.S., O.S. and T.S. contributed to immunogen design. S.B. and J.K.D. performed, and J.R.Y. and J.C.P. supervised mass spectrometry glycan profiling. P.J.M. and M.S. performed, and S.C. supervised NHP B cell FACS analysis. S.W. and A.G. contributed, and G.O. and A.B.W. supervised cryo-EM imaging, data processing and model building. D.L. performed nanodisc assembly reactions. P.K. and S.T. performed mouse immunizations. D.L., E.G., R.T., S.E., N.A., D.G. and M.K. produced purified proteins. W-H.L. contributed to cryo-EM sample preparation. S.H. provided mRNA immunogens. T.S., A.B.W. and W.R.S. supervised the study. G.O., T.S., A.B.W. and W.R.S. edited the manuscript. All authors reviewed the manuscript.

