## Supplementary information for "Virus glycoprotein nanodisc platform for vaccine design"

Supplementary figures 1 – 4

Supplementary table 1

Supplementary video 1 legend

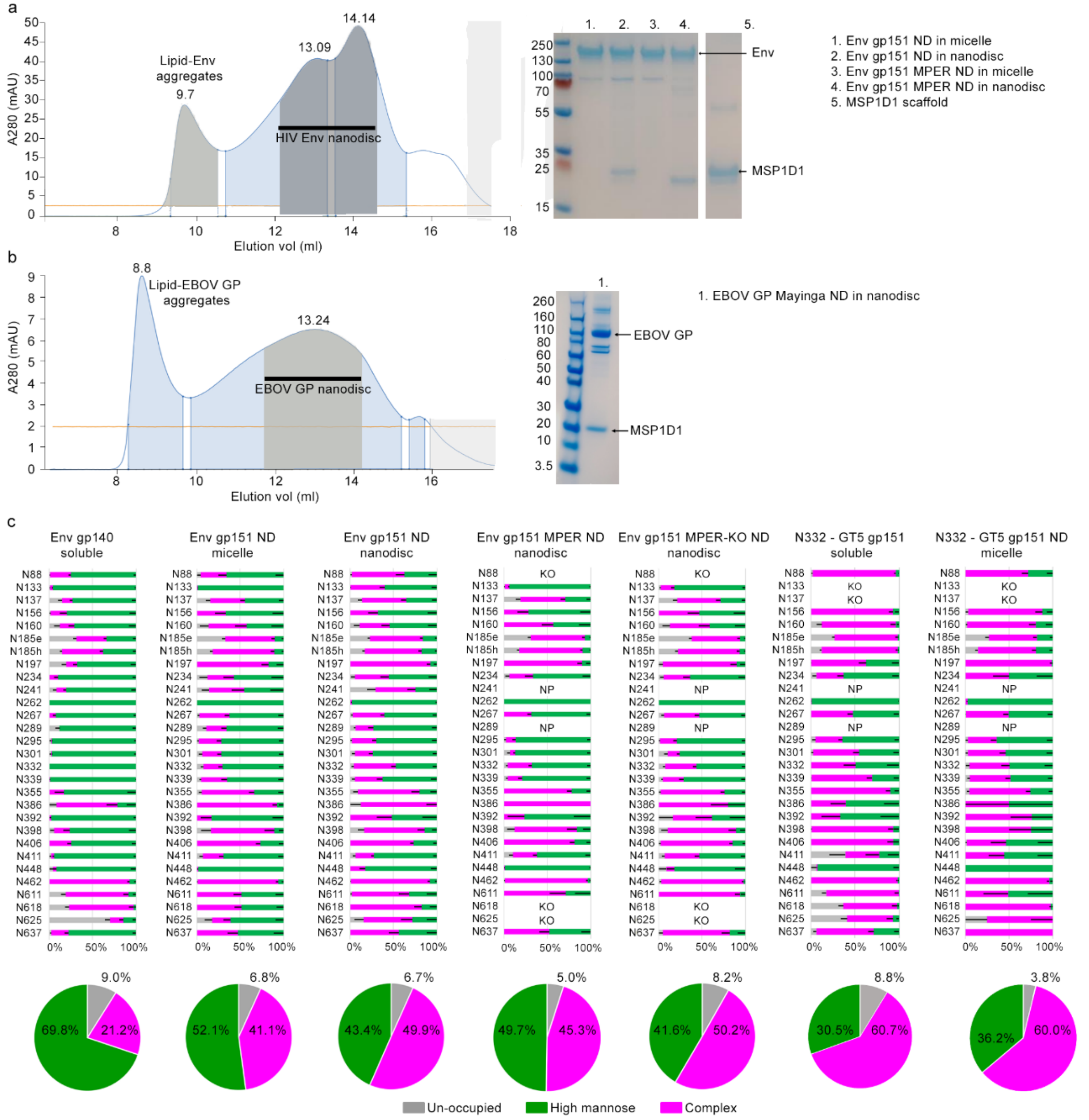


**Supplementary figure 1** Quality control steps of nanodiscs. a-b) Typical size exclusion chromatogram (left) showing wide peaks indicative of glycan heterogeneity and GP flexibility on nanodisc also seen on nsEM controls. SDS-PAGE analysis (right) confirms purity of the samples and expected bands corresponding the glycoprotein monomer with added mass from glycans, as well as the presence of the scaffold protein. c) Comprehensive glycotype analysis of HIV glycoproteins are shown with error bars showing standard error of mean. Data confirms the presence of all expected glycan sites and shows higher proportion of complex type glycans in transmembrane Env samples, including the nanodisc, that were tested for MPER targeting immunogen development, and high proportion of complex type glycans for N332–GT5 soluble version of the immunogen, closely matching that of the transmembrane version.

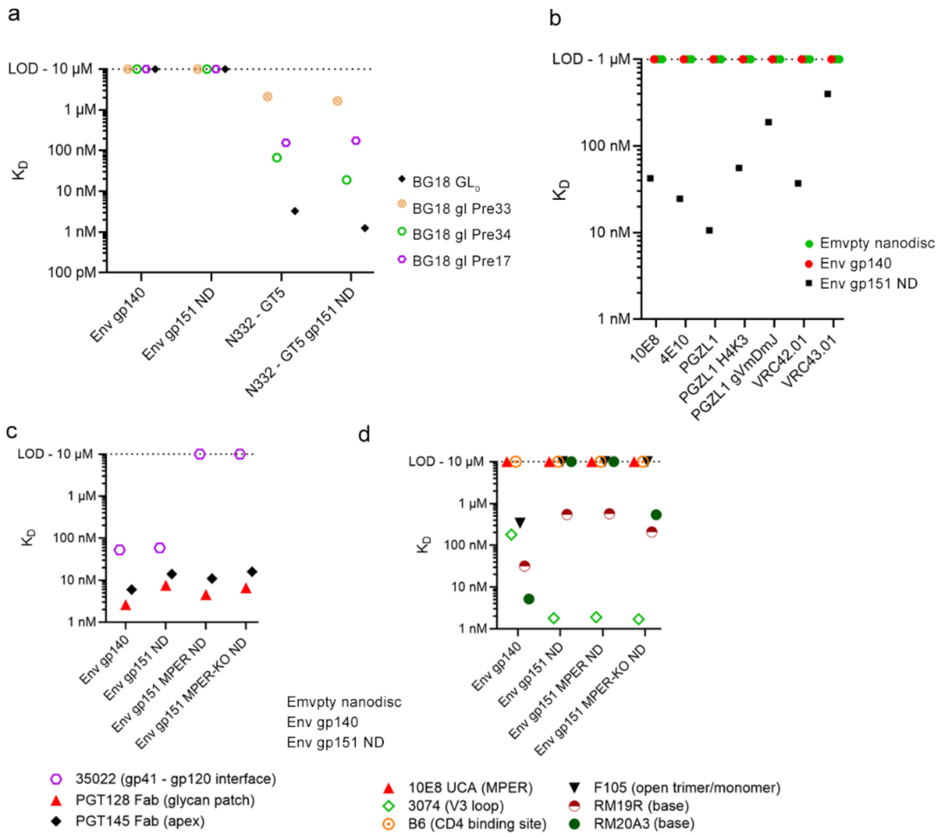


**Supplementary figure 2** Additional SPR data. a) A transmembrane version of germline targeting vaccine construct N332 - GT5 was assembled into nanodisc using the ND assembly workflow. N332-GT5 gp151 ND showed comparable affinities between the soluble and transmembrane version of the immunogen against human germline antibodies. Affinities were measured using SPR modality A (capture nanodisc, flow Fab). b) Affinities of a panel of HIV Env MPER targeting antibodies to Env gp151 ND measured with SPR modality B (capture IgG, Flow nanodisc). Empty nanodiscs served as controls to assess direct antibody-lipid interactions. While MPER-targeting antibodies have previously been described to interact with lipids, no binding to empty nanodiscs was detected at 1μM analyte concentration. c) Antigenic profiling of Env constructs used in pilot MPER targeting immunogen development with neutralizing and (d) non-neutralizing antibodies as measured by SPR modality A.

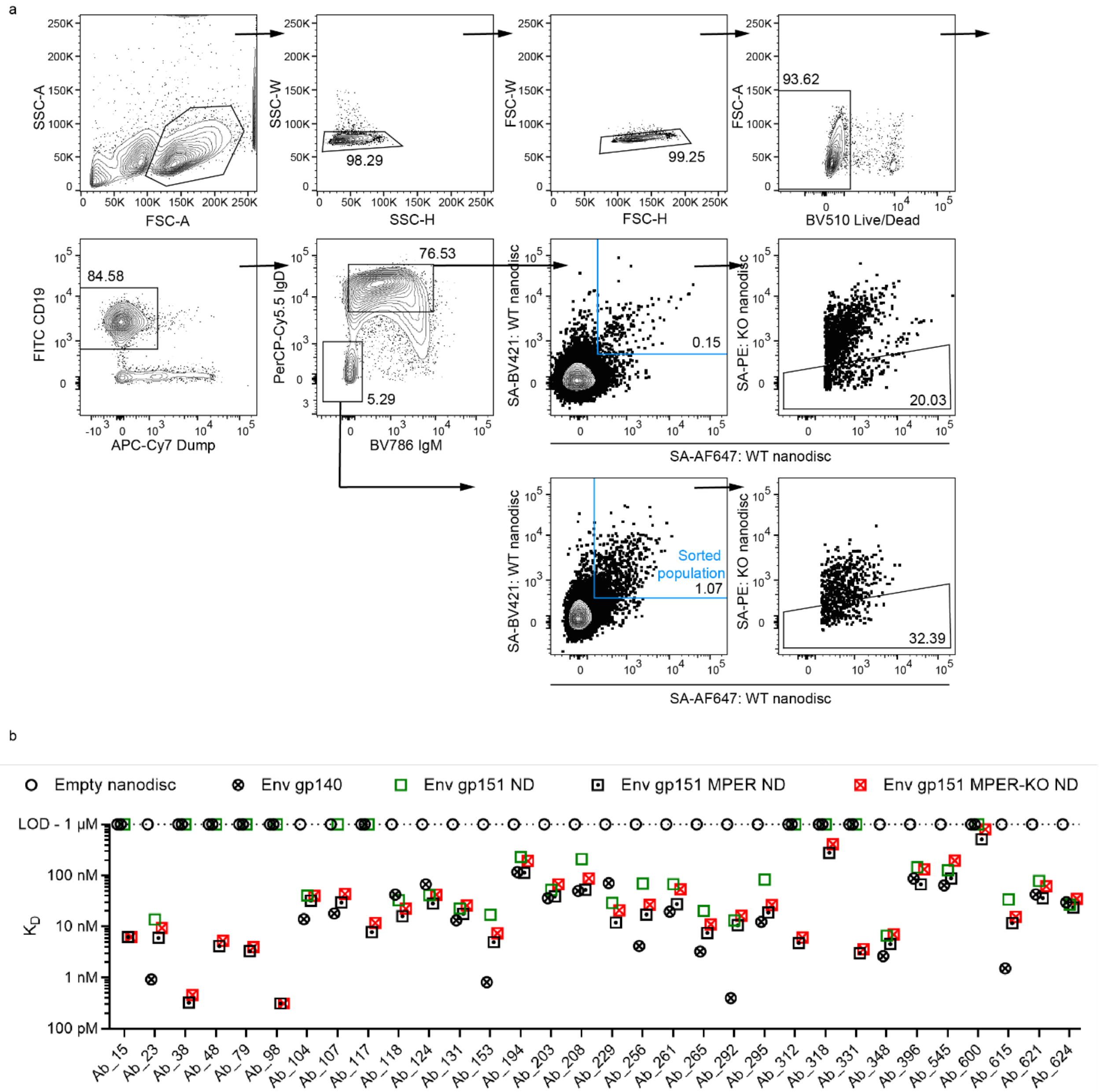


**Supplementary figure 3** Additional analysis of mouse model data. a) Complete mouse B cell gating strategy in reference to Fig 4. b) Affinities of sorted, sequenced and purified monoclonal antibodies from Fig 4d that showed binding to at least one of the Envs as measured by SPR modality B.

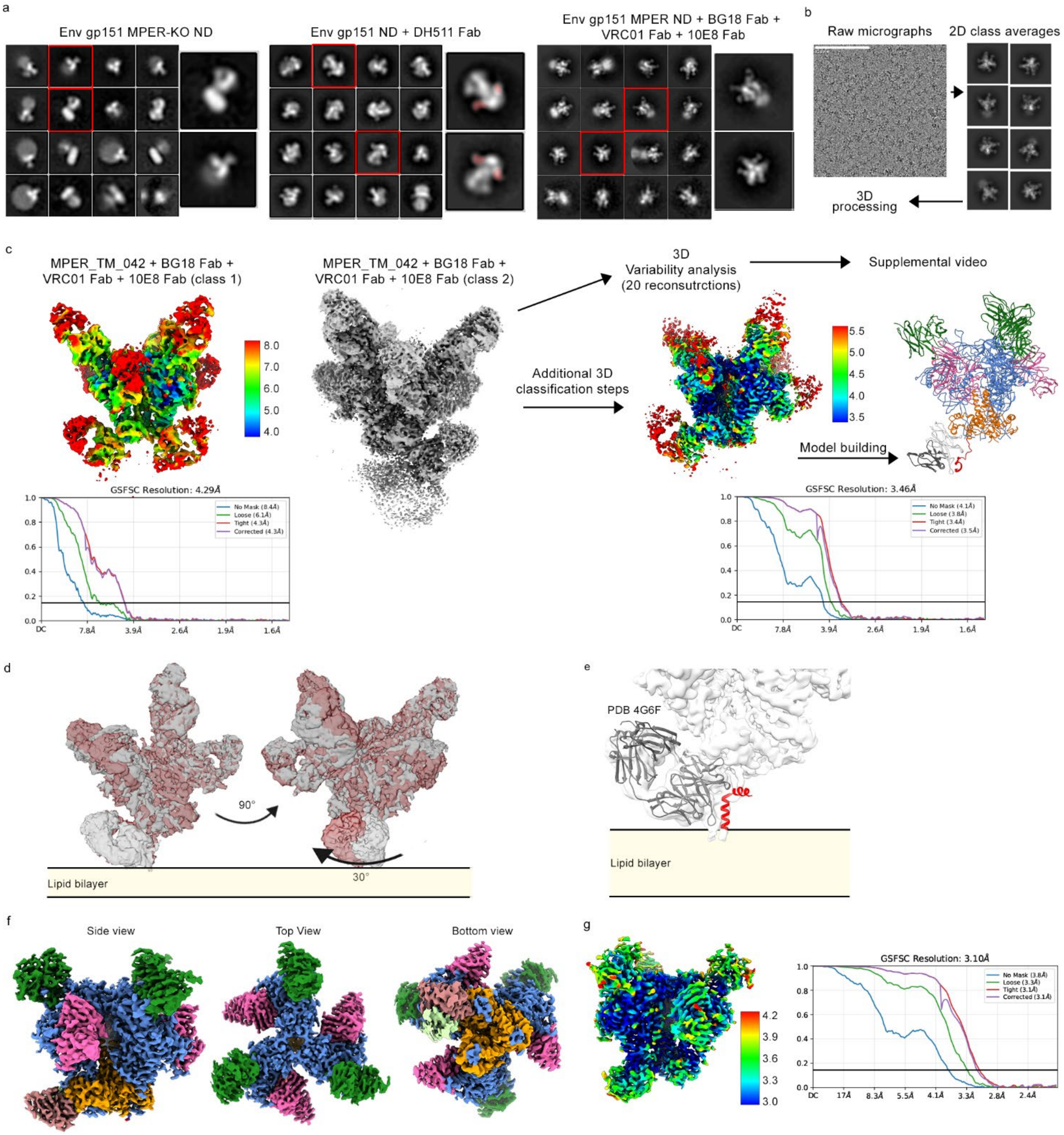


**Supplementary figure 4** EM analysis of glycoprotein nanodiscs. a) Representative quality control 2D class averages from negative stain EM imaging showing unliganded nanodiscs with typical features of 10 nm disc at the base of the particle and trimeric feature of the glycoprotein. MPER targeting Fab DH511 (highlighted in red) is visible at the base of the trimer density. Env gp151 MPER ND nanodisc was complexed with BG18, VRC01 and 10E8 Fabs and presences of all three Fabs was confirmed by 2D class averages. b) The latter complex was subjected to structural studies by cryo-EM. c) All data was processed within cryosparc software suite using various processing workflows leading to final maps with one or two copies of 10E8 bound. Local resolution maps and overall GSFSC resolution estimates are given for each final map. The largest number of particles had one 10E8 bound. This particle class then classified into distinct conformations using 3D variability analysis to visualize flexibility in the 10E8 epitope. Further 3D classification led to highest resolution reconstruction at 3.5 Å, which was used for model building. d) Two 3D refinements (grey and red) overlaid from the end-points of 3D variability analysis showing ~30° lateral twist of the 10E8 antibody. e) X-ray structure of 10E8 (grey ribbon) in complex with MPER peptide (red; PDB 4G6F) fitted into lowpass filtered cryo-EM map with single 10E8 bound. f) Cryo-EM map of Env gp140 (gp120 in blue, gp41 in orange) in complex with BG18 (green), VRC01 (pink), and 3BC315 (HC in salmon, LC in light green). g) Local resolution and GSFCS resolution estimation of the Env gp140 cryo-EM map.

**Supplementary table 1** Cryo-EM data collection, refinement and validation statistics

| | Env gp151 MPER ND + BG18 Fab + VRC01 Fab + 10E8 Fab (class 1) (EMD-70471) (PDB 9OGM) | Env gp151 MPER ND + BG18 Fab + VRC01 Fab + 10E8 Fab (class 2) (EMD-70470) | BG505 MD39.3 gp140 + BG18 Fab + VRC01 Fab + 3BC315 Fab (EMD-70469) (PDB 9OGL) |
|---|---|---|---|
| **Data collection and processing** | | | |
| Microscope | TFS Glacios | TFS Glacios | TFS Glacios |
| Voltage (keV) | 200 | 200 | 200 |
| Camera | TFS Falcon 4i | TFS Falcon 4i | TFS Falcon 4i |
| Collection mode | Counting | Counting | Counting |
| Magnification | 190,000x | 190,000x | 190,000x |
| Pixel size at detector (Å) | 0.718 | 0.718 | 0.718 |
| Total electron exposure (e–/Å$^2$) | 44 | 44 | 45 |
| Exposure rate (e-/pixel/sec) | 8.93 | 8.93 | 4.98 |
| Number of EER frames | 40 | 40 | 40 |
| Defocus range (μm) | 0.8 to -2.0 | 0.8 to -2.0 | 0.8 to -2.0 |
| Automation software | EPU | EPU | EPU |
| Micrographs collected (no.) | 12,067 | 12,067 | 8,010 |
| Micrographs used (no.) | 11,059 | 11,059 | 7,427 |
| Initial particle images (no.) | 306,760 | 306,760 | 886,907 |
| Final particle images (no.) | 63,366 | 7,964 | 243,503 |
| Symmetry | C1 | C1 | C3 |
| Map pixel size (Å) | 0.718 | 0.718 | 1.034 |
| Map resolution (masked/unmasked Å) | 3.5/4.1 | 4.3/8.4 | 3.1/3.8 |
| FSC threshold | 0.143 | 0.143 | 0.143 |
| Map sharpening *B* factor (Å$^2$) | -61.3 | -13.8 | -68.3 |
| Map resolution range (Å) | 3.0-6.0 | 4.0-8.0 | 3.0-4.2 |
| **Refinement** | | | |
| Initial model used (PDB code) | 3NGB, 6DFG, 4G6F | n/a | 5CCK, 6DFG, 3NGB |
| Refinement package | Phenix real space refine | n/a | Phenix real space refine |
| Model resolution (Å) | 3.8 | n/a | |
| FSC threshold | 0.5 | n/a | 0.5 |
| EMRinger score | 2.85 | n/a | 2.79 |
| CC (mask) | 0.81 | n/a | 0.78 |
| *Model composition* | | | |
| Non-hydrogen atoms | 27,059 | n/a | 27,288 |
| Protein residues | 3,258 | n/a | 3,262 |
| Ligands | 115 | n/a | 133 |
| *Mean B factors (Å$^2$)* | | | |
| Protein | 56.16 | n/a | 71.57 |
| Ligand | 60.74 | n/a | 85.67 |
| *R.m.s. deviations* | | | |
| Bond lengths (Å) | 0.006 | n/a | 0.006 |
| Bond angles (°) | 0.977 | n/a | 0.999 |
| *Validation* | | | |
| MolProbity score | 1.65 | n/a | 1.04 |
| Clashscore | 5.62 | n/a | 1.77 |
| Poor rotamers (%) | 0.74 | n/a | 0.46 |
| *Ramachandran plot* | | | |
| Favored (%) | 95.05 | n/a | 97.50 |
| Allowed (%) | 4.95 | n/a | 2.50 |
| Disallowed (%) | 0.00 | n/a | 0.00 |
| Cβ outliers (%) | 0.00 | n/a | 0.00 |
| CaBLAM outliers (%) | 3.49 | n/a | 2.69 |

**Supplementary video 1.** Structural flexibility in the MPER epitope. 3D Variability analysis was performed for Env gp151 MPER ND + BG18 Fab + VRC01 Fab + 10E8 Fab (class 1) cryo-EM particle set. Interpolating between 20 3D reconstructions show ~30° lateral movement of 10E8 Fab in relation to ectodomain and an associated ectodomain displacement and MPER domain remodeling.
